# Cell-envelope synthesis is required for surface-to-mass coupling, which determines dry-mass density in *Bacillus subtilis*

**DOI:** 10.1101/2021.05.05.442853

**Authors:** Yuki Kitahara, Enno R. Oldewurtel, Sean Wilson, Yingjie Sun, Ethan C. Garner, Sven van Teeffelen

## Abstract

Cells must increase their volumes in response to biomass growth to maintain intracellular mass density, the ratio of dry mass to cell volume, within physiologically permissive bounds. To increase volume, bacteria enzymatically expand their cell envelopes and insert new envelope material. Recently, we demonstrated that the Gram-negative bacterium *Escherichia coli* expands cell-surface area rather than volume in proportion to mass. Here, we investigate the regulation of cell-volume growth in the evolutionarily distant *Bacillus subtilis*. First, we demonstrate that the coupling of surface growth to mass growth is conserved in *B. subtilis*. Therefore, mass density changes with cell shape at the single-cell level. Interestingly, mass density varies by more than 30% when we systematically change cell width by modulation of cell-wall insertion, without any effect on mass-growth rate. Second, we demonstrate that the coupling of surface- and mass growth is broken if peptidoglycan or membrane synthesis are inhibited. Once transient perturbations are relieved, the surface-to-mass ratio is rapidly restored. In conclusion, we demonstrate that surface-to-mass coupling is a conserved principle for volume regulation in bacteria, and that envelope synthesis provides an important link between surface growth and biomass growth in *B. subtilis*.

## Introduction

In bacteria and other organisms, the cytoplasm is crowded with macromolecules, notably protein, RNA, and DNA, which occupy about 30-40% of the volume (***Cayley et al., 1991**; **Zimmerman and Trach, 1991***). Cytoplasmic crowding is important for cell physiology as it directly impacts macromolecular diffusion (***Konopka et al., 2009**; **Delarue et al., 2018***), molecular interactions (***Zhou et al., 2008***), and chromosome organization (***Yang et al., 2020***). Furthermore, it was also suggested that crowding is maximizes biomass growth rate (***Dourado and Lercher, 2020**; **Vazquez, 2010***). To maintain the density of macromolecules and other cytoplasmic components within physiologically permissive bounds or to achieve optimal crowding cells must coordinate their volume growth rate with the rate of biomass growth.

We recently studied this problem in the Gram-negative bacterium *Escherichia coli **(Oldewurtel et al., 2019***). By measuring single-cell dry mass and cell dimensions independently using quantitative phase microscopy, we showed that cells control cell-volume growth indirectly on the timescale of about one generation: Cells increase their surface area rather than volume in proportion to dry-mass growth. Thus, they maintain a constant ratio of cell-surface area ***S*** to total cellular dry mass ***M***.

During steady-state growth, when cell width remains almost constant, this coupling guarantees that cell volume grows roughly in proportion to mass, because surface area, volume, and length increase approximately in proportion to one another. However, if cells systematically increase their width, for example after a nutrient upshift, cell volume grows faster than surface area. Thus, the cytoplasm is diluted, and dry-mass density drops (***Oldewurtel et al., 2019***).

The robust coupling of surface area and dry mass in *E. coli* implies that surface area increases by the same relative amount as dry mass, independently of dry-mass density, turgor pressure, or instantaneous growth rate. The surface-to-mass coupling might thus be metabolic in nature, through the production of a rate-limiting cell-envelope component, while other physiological processes such as crowding and turgor pressure have no apparent influence on surface growth on short timescales, in agreement with previous work by ***Rojas et al.** (**2014***). However, the metabolic pathways responsible for surface-to-mass coupling remain to be identified in *E. coli* or any other bacterium.

Here, we investigate how the Gram-positive bacterium *Bacillus subtilis* coordinates volume and biomass growth. Gram-negative and Gram-positive bacteria differ in envelope structure and envelope growth in fundamental ways: Gram-negative bacteria are surrounded by a thin peptidoglycan cell wall and by a mechanically important outer membrane. On the contrary, Gram-positive bacteria lack an outer membrane but are surrounded by a much thicker cell wall. Furthermore, osmotic pressure (turgor) has an influential role in surface-area expansion in *B. subtilis* but not in *E. coli*. More specifically, *B. subtilis* changes its rate of surface growth in response to changes of turgor pressure (***Rojas et al., 2017***), while *E. coli* does not (***Rojas et al., 2014***). It thus remains unclear whether the robust surface-to-mass coupling observed in *E. coli (**Oldewurtel et al., 2019***) is maintained in *B. subtilis*. Furthermore, the role of the insertion of peptidoglycan and other envelope components for surface growth remains to be explored.

Using quantitative phase microscopy, we demonstrate here that surface and mass are robustly coupled during growth of *B. subtilis*, even if cell width and therefore dry-mass density changes. Specifically, dry-mass density is inversely proportional to width at the single-cell level. Furthermore, we observed similar correlations at the population level when systematically varying cell width by modulating the expression of class-A penicillin binding proteins (aPBPs) as previously described (***Dion et al., 2019***). Upon increase of cell width, dry-mass-density decreases by up to 30%, but biomass growth rate and cell-wall insertion remain remarkably constant. Thus, dry-mass density and crowding do not govern cell-envelope growth.

Which pathway is responsible for the coupling between surface and dry-mass growth? Physically, cell-surface area is governed by the peptidoglycan cell wall. Thus, cell-wall cleaving autolysins are strictly required for growth. In visionary and influential work, ***Koch** (**1983***) suggested that ‘smart autolysins, are activated based on mechanical stress in the cell wall, which, in turn, is caused by turgor pressure. However, more recent works imply that the MreB-linked cell-wall insertion machinery provides the major regulator of cell elongation in *B. subtilis (**Daniel and Errington, 2003**; **Billaudeau et al., 2017**; **Domínguez-Cuevas et al., 2013**; **Rojas et al., 2017**; **Sun and Garner, 2020***). A regulatory role of peptidoglycan insertion for autolytic activity is supported by previous studies suggesting that the two redundantly essential cell-wall hydrolases of *B. subtilis*, LytE and CwlO, are controlled by the three MreB homologs (***Carballido-López et al., 2006**; **Domínguez-Cuevas et al., 2013***). Furthermore, the amount of moving MreB filaments and cell-envelope growth are highly correlated across different growth conditions (***Sun and Garner, 2020***), which is compatible with a rate-limiting role of MreB-based cell-wall insertion. However, a molecular mechanism linking cell-wall insertion and cell-wall expansion has not been identified. Furthermore, there is also evidence against this hypothesis: Specifically, sub-lethal concentrations of cell-wall antibiotics such as fosfomycin do not affect cell-elongation rate (***Rojas et al., 2017***). Furthermore, we recently discovered that peptidoglycan insertion is not rate-limiting in *E. coli (**Oldewurtel et al., 2019***), contrary to wide-spread belief (***Höltje, 1998***). Thus, the connection between cell-wall insertion, biomass growth, and surface expansion remains unclear.

Another essential envelope component is the cytoplasmic membrane. Previously, ***Rojas et al.** (**2017***) provided combined experimental and model-based evidence that membrane tension is important to facilitate cell-wall insertion, which, together with turgor pressure, might be responsible for driving cell-wall expansion (***Rojas et al., 2017***). Furthermore, ***Müller et al.** (**2016***) and ***Zielińska et al. (2020***) demonstrated that membrane fluidity and membrane micro-domain organization affects cell-wall insertion. Interestingly, inhibition of membrane synthesis through glycerol starvation in a glycerol auxotroph increases buoyant mass density (***Mindich, 1970***), which is compatible with the idea that surface area increases more slowly than mass during the arrest of membrane synthesis. Whether membrane insertion constitutes a direct link between cell-surface area and biomass growth remains to be investigated.

In this work we demonstrate that both cell-wall insertion and cell-membrane insertion are required for proper surface growth and for the maintenance of ***S***/***M***. If either of the two processes is inhibited, surface growth is severely reduced, while biomass growth continues. Interestingly, though, cell-wall insertion is not directly coupled to cell-surface growth. Instead, we observe a delay between the arrest of peptidoglycan insertion and the reduction of surface growth, in agreement with previous observations (***Misra et al., 2013***). Subsequently, surface growth continues at a reduced rate, even though cell-wall insertion is inhibited. Similarly, cells can reduce surface growth even if the rate of peptidoglycan insertion remains high. Thus, cell-wall insertion is important but not rate-determining for cell-surface growth. Similarly, we find that membrane insertion is required for proper surface growth but the visible overproduction of membrane does not lead to increased surface growth. Once the perturbation of envelope synthesis is relieved, ***S**/**M*** returns rapidly to its pre-treatment value.

Together, our experiments demonstrate that cells control cell-volume growth indirectly, by coupling surface growth to mass growth, with an important role of envelope synthesis for cell-surface growth.

## Results

### Cells grow surface rather than volume in proportion to biomass

To study the relationship between cell-volume growth and biomass growth in *B. subtilis*, we quantified single-cell dimensions and dry mass of live cells in absolute terms using quantitative phase microscopy, similarly to our recent measurements on *E. coli* (***Oldewurtel et al., 2019***). However, because the cell wall is a thick and uneven layer in Gram-positive bacteria, we decided to measure cytoplasmic properties rather than whole-cell properties. Specifically, we calculated cytoplasmic volume ***V***, surface area ***S***, and width ***W*** based on two-dimensional cell contours from phase-contrast images using the Morphometrics tool (***Ursell et al., 2017***), after calibration based on the membrane dye FM 4-64 (Figure 1 - figure supplement 1). To obtain cytoplasmic mass, we first measured the total cellular dry mass ***M***_tot_ using Spatial Light Interference Microscopy (SLIM) (***Wang et al., 2011***), a variant of quantitative phase microscopy. This measurement is accurate and precise, as demonstrated in *E. coli*. Subsequently, to obtain cytoplasmic mass ***M***, we subtracted a constant fraction of 14%, which represents the dry mass of the cell wall obtained by bulk measurements (two biological replicates: 13.8%, 14.2%). Other extra-cytoplasmic contributions to total mass, for example from periplasmic proteins, are significantly smaller than the cell-wall contribution and thus implicitly allocated to the cytoplasmic mass for simplicity (see Materials and Methods).

First, we grew cells to exponential phase in different growth media in batch and took snapshots on agarose pads (Figure 1A). Despite almost threefold differences in average growth rate, the average cytoplasmic dry-mass density of about 0.31 – 0.33 g/ml (Figure 1B) varies by no more than 5% between conditions. According to our overestimation of cytoplasmic mass by about 5%, absolute values of cytoplasmic mass density might be slightly lower (0.29 – 0.31 g/ml). These results are in agreement with independent measurements of average refractive index through immersive refractometry (Figure 1 - figure supplement 2). Interestingly, the mass density of *B. subtilis* is very similar to mass densities recently measured in *E. coli (**Oldewurtel et al., 2019***).

**Figure 1.**
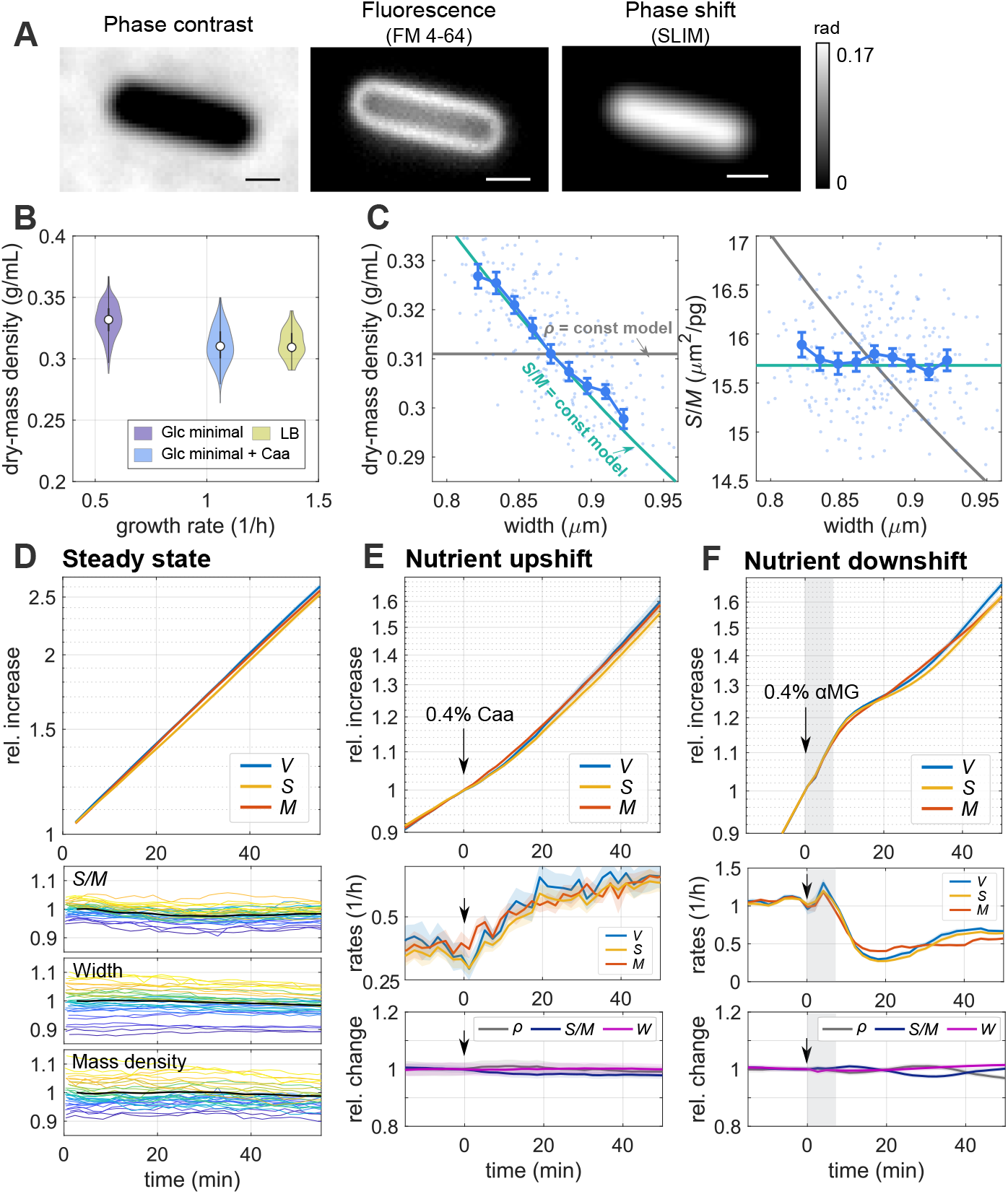
*B. subtils* controls volume indirectly by growing surface in proportion to dry mass. **A:** Snapshots of a wild-type cell (PY79) labeled with the membrane dye FM4-64, in minimal medium with glucose and casamino acids (S7_50_+GlcCaa) taken by phase-contrast microscopy, fluorescence microscopy, and SLIM (grayscale bar: phase shift). Scale bars 1 μm. **B:** Distribution of single-cell dry-mass density of wild-type cells grown to exponential phase in minimal medium with glucose (S7_50_+Glc), glucose and casamino acids (S7_50_+GlcCaa), and in LB (white circles = median; grey rectangles = interquartile range). See Figure 1 - figure supplement 3 for other properties. **C:** Width dependency of dry-mass density and surface-to-mass ratio in S7_50_+GlcCaa medium (dots: single-cell measurements, blue lines: binned averages ± SE, green lines: model prediction for spherocylinder with constant surface-to-mass ratio, gray lines: model prediction for spherocylinder with constant dry-mass density). **D:** Single-cell time lapse of filamentous cells (bAB56) on agarose pad (S7_50_+GlcCaa). To avoid cell division, MciZ was induced 30 min prior to microscopy. Relative increase (top) of volume, surface and dry mass (Solid lines + shadings = average ± SE). Bottom: Relative change of single-cell width, surface-to-mass ratio and dry-mass density (black lines: average values). **E:** Single-cell time lapse of filamentous cells (bAB56) on agarose pad during nutrient upshift from S7_50_+Glc to S7_50_+GlcCaa at time = 0 min (by addition of casamino acids to the top of the agarose pad). To avoid cell division, MciZ was induced 50 min prior to microscopy. Relative increase (top) and rates (mid) of volume, surface and dry mass. Bottom: Relative change of average dry-mass density, surface-to-mass ratio, and width (Solid lines + shadings = average ± SE). **F:** Single-cell time lapse of filamenting cells (bAB56) during nutrient downshift from S7_50_+GlcCaa to S7_50_+GlcCaa + glucose analogue (alpha methylglucoside). To avoid cell division, MciZ was induced 30 min prior to microscopy. Shaded background indicates region of droplet addition that causes transient measurement distortions. Otherwise the same as in E.

Similar to *E. coli*, single-cell variations in dry-mass density are also remarkably small (**CV** ≈ 3-5%). However, we note that about 80% of cells imaged here contain a septum (complete or non-complete) according to membrane staining, and thus possibly represent two cells separated by a membrane. Thus, single-cell variations might be slightly underestimated.

The narrow distribution of dry-mass density could come about in either of two ways: Cells could control dry-mass density directly, by increasing cell volume in proportion to dry mass, or cells could control dry-mass density indirectly, for example by increasing surface area rather than volume in proportion to dry mass, as recently observed in *E. coli (**Oldewurtel et al., 2019***). Drymass density ***ρ*** = ***M**/**V*** can be expanded as the ratio of surface-to-volume and surface-to-mass ratios, ***ρ*** = (***S**/**V***)/(***S**/**M***). For spherocylinders such as *B. subtilis* or *E. coli*, ***S**/**V*** scales approximately inversely with width ***W*** according to ***S**/**V*** ≈ 4/***W***. We can thus write ***ρ*** ≈ 4/[***W***(***S***/***M***)]. If cells grew surface area in proportion to mass, independently of any change of dry-mass density, we would expect that dry-mass density is inversely correlated with cell width, while the surface-to-mass ratio ***S**/**M*** shows no or weak dependency on width. Alternatively, if cells control volume in direct response to mass growth, we would expect that ***S**/**M*** and width are negatively correlated.

To distinguish the two possibilities we studied correlations between single-cell values of dry-mass density with width and of ***S**/**M*** with width (Figure 1C), where every point represents a single cell. We found that ***ρ*** shows an inverse proportionality with width (Figure 1C) while ***S**/**M*** shows hardly any dependency on width. This behavior is also seen in a different growth medium (Figure 1 - figure supplement 4). Here and in the following we consider the ratios of cytoplasmic surface area and mass. However, the same relation holds, if we were to normalize with respect to total mass ***M***_tot_, simply because ***M*** is calculated as a constant fraction of ***M***_tot_.

Our observations thus suggest that *B. subtilis* controls cell volume indirectly, by growing surface area rather than volume in proportion to mass,just like the Gram-negative *E. coli* (***Oldewurtel et al., 2019***).

To study whether the surface-to-mass ratio is maintained on long timescales, we also conducted time-lapse microscopy experiments. To study cells for more than one generation of growth, we inhibited cell division by inducing the expression of MciZ, a peptide that inhibits Z-ring formation (***Handler et al., 2008***), 30-50 min prior to microscopy, using the strain bAB56 (*mciZ*::spec-pHyperSpank-*mciZ*), similar to previous work (***Dion et al., 2019***) and as specified in Supplementary File 1. Remarkably, single-cell values of ***S**/**M*** and width remain nearly constant during one mass doubling (Figure 1D). Accordingly, mass density remains also nearly constant on this timescale (Figure 1D). These slow temporal variations are reflected by slowly decaying temporal auto-correlation functions (ACF) of all three quantities (Figure 1 - figure supplement 5A) and our observations are consistent with tight correlations between the rates of surface and mass growth (Figure 1 - figure supplement 5B).

### Surface expansion and mass growth are robustly coupled during nutrient shifts

In *E. coli* we previously observed that the surface-to-mass ratio is maintained nearly constant during rapid changes of growth rate, apart from transient variations ascribed to changes of turgor pressure (***Oldewurtel et al., 2019***). Those observations gave rise to a new growth law: The instantaneous rate of surface growth is directly proportional to biomass growth d*S*/d*t* = *α*d**M**/d*t*, where *α* remains constant or changes slowly on the generation time scale. To test whether this relation also holds in *B. subtilis*, we studied single cells in time-lapse microscopy experiments on agarose pads during nutrient shifts (Figure 1E,F, Figure 1 - video 1, Figure 1 - video 2).

For a nutrient upshift we grew cells in minimal medium supplemented with glucose (S7_50_+Glc) and then added casamino acids 30 min after starting the experiment (Figure 1E). Despite the 60% increase of biomass growth rate over the course of about 20 min, cells increase mass and surface area synchronously, thus maintaining ***S**/**M*** robustly constant, both in terms of the population average (Figure 1E) and at the single-cell level (Figure 1 - figure supplement 6). Since width remains almost constant, mass density also remains nearly constant.

We also measured the behavior during a nutrient downshift. To that end we exposed cells growing on minimal medium supplemented with glucose and casamino acids (S7_50_+GlcCaa) to 0.4% alpha methylglucoside (alpha-MG), a non-metabolisable analogue of glucose (***Freese et al., 1970***). Upon alpha-MG addition growth rates of mass, surface, and volume drop almost synchronously by more than two-fold within 10 min (Figure 1F). Surface and volume growth rates undershoot slightly and then oscillate around the constant mass growth rate, which leads to small deterministic variations of ***S**/**M*** and mass density (of about 2%), also observed at the single-cell level (Figure 1 - figure supplement 6). Similar to the nutrient upshift, cell width remains almost constant.

In conclusion, *B. subtilis* adjusts the rate of surface growth rapidly during nutrient upshift and downshift, and 〈***S**/**M***〉 remains almost constant, suggesting that the growth law identified in *E. coli* is approximately conserved in *B. subtilis*.

### Modulation of cell width through class-A PBPs changes average dry-mass density without perturbing growth rate

During steady-state growth conditions we observed that dry-mass density changed with increasing cell width. Here, we aimed to test whether this is also true if cell width changes on average. Different from *E. coli, B. subtilis* does not show pronounced changes of average cell width between different growth media (Figure 1 - figure supplement 3). We therefore aimed to change cell width genetically.

It was previously described that cell width changes in response to the balance between the activities of two different peptidoglycan-synthesizing machineries, the MreB-actin-rod complex and the class A penicillin-binding proteins (aPBPs) (***Dion et al., 2019***). We thus modulated the expression level of the major aPBP PonA expressed from an IPTG-inducible promoter using strain bMD586 (*yhdG*::*cat-pHyperSpank-ponA*,*ponA*::*kan*), or we used mutants that lack either PonA(Δ*ponA*;bKY42) or all four known aPBPs (Δ4; bSW164). As expected from ***Dion et al.** (**2019***), cell width of bMD586 increased continuously with increasing IPTG levels, while width was reduced in the *ΔponA* and Δ4 strains with respect to the wildtype (Figure 2A).

**Figure 2.**
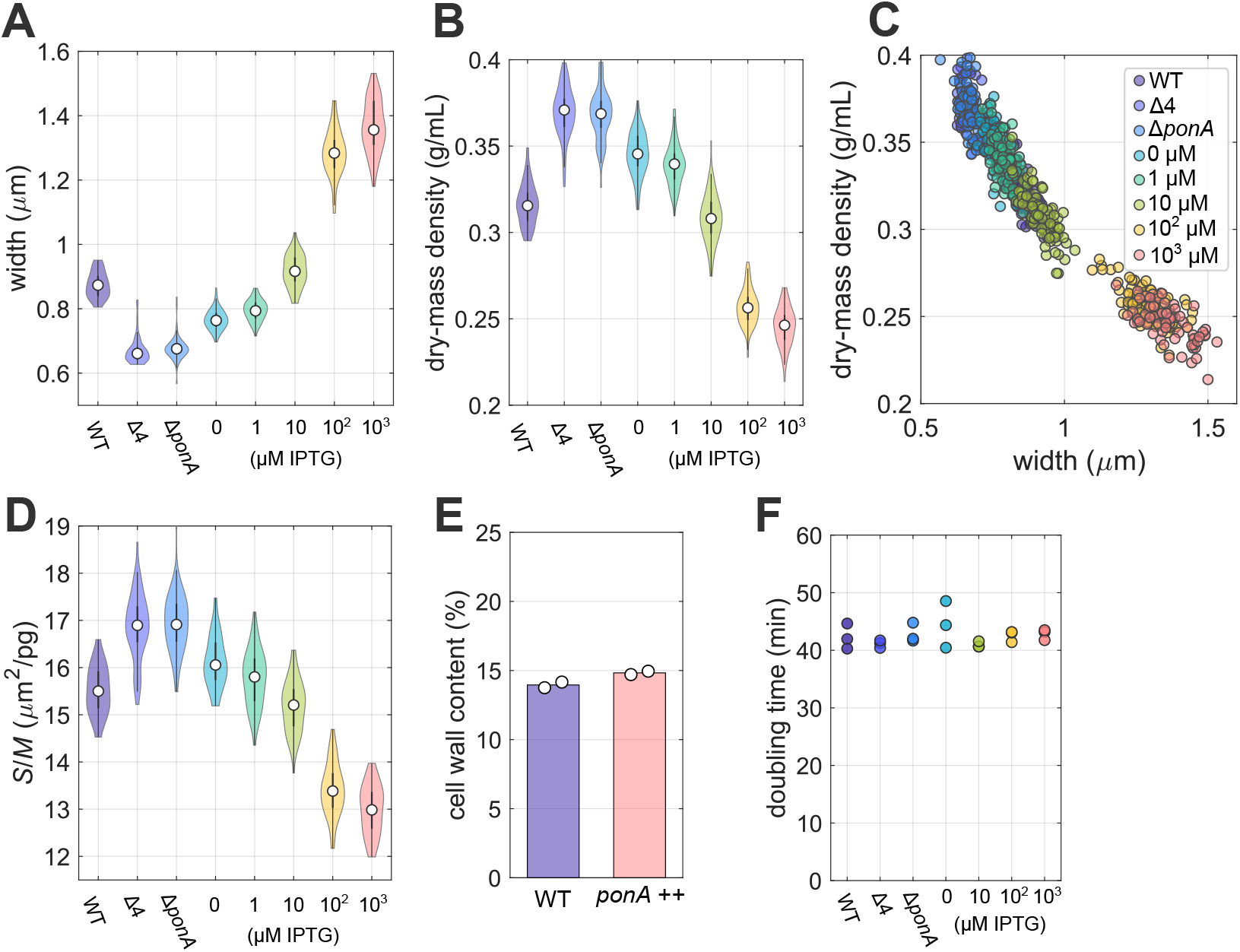
Systematic modulation of cell width leads to changes of average dry-mass density without perturbing mass growth rate. **A-D:** Width (**A**), dry-mass density (**B-C**), and surface-to-mass ratio (**D**) under different expression levels of aPBPs. Snapshots of wild-type, bSW164, bKY42, and bMD586 cells during steady-state growth in S7_50_+GlcCaa medium. To titrate PonA expression level in bMD586, IPTG was added from 1 to 1000 μM. A,B,D: Violin plots with median (white circles) and interquartile range (grey rectangles), C: Single-cell data. **E:** Cell-wall content per cellular dry mass of wild-type and bMD586 (1mM IPTG) cells grown in S7_50_+GlcCaa medium (white circles = biological replicates) **F:** Bulk doubling times of wildtype, bSW164, bKY42, and bMD586 in S7_50_+GlcCaa medium. The doubling times were calculated by fitting values of optical density between 0.02 to 0.3 (circles = biological replicates).

As hypothesized, dry-mass density decreases with increasing average cell width (Figure 2B). Inverse correlations between density and width are also observed at the single-cell level (Figure 2C). In comparison to the wildtype, mass density decreases or increases by about 20% on average upon overexpression or depletion of aPBPs, respectively. We confirmed these changes through an alternative technique, immersive refractometry (Figure 2 - figure supplement 1).

Upon modulation of aPBP levels we also observed a decrease of the surface-to-mass ratio with increasing aPBP expression (Figure 2D). Mass density is approximately inversely proportional to the product of both width and ***S**/**M*** (***ρ*** ≈ 4/***W***/(***S**/**M***)). Since relative variations in 〈***S**/**M***〉 are about three-fold smaller than relative variations in width, changes in mass density (Figure 2B,C) are dominated by changes of width.

Previously, ***Dion et al.*** (***2019***) showed that the cell wall is thicker upon high PonA expression than in wild-type cells, suggesting that the amount of cell-wall material per surface area is increased. We thus speculated that the decrease of 〈**S**/**M**〉 might be a consequence of the thicker cell wall, while the rate of cell-wall insertion per mass and thus the amount of cell-wall material per mass remains unchanged. Indeed, we observed that the total amount of cell-wall material per biomass remains constant upon PonA overexpression (Figure 2E). The reduction of ***S**/**M*** is thus consistent with the previously observed increase of cell-wall thickness.

Previous theoretical work suggests that mass growth rate might depend on intracellular density and crowding (***Vazquez, 2010**; **Dourado and Lercher, 2020***). However, we found that growth rate remains constant across the wide range of mass densities explored here (Figure 2F; for growth curves see Figure 2 - figure supplement 2). Previously, (***Popham and Setlow, 1996***) reported reduced growth rates of mutants lacking individual or all aPBPs grown at 37°C. However, this reduction can be attributed to increased cell lysis (***Meeske et al., 2016**; **Dion et al., 2019***), while growth rate remains high at the single-cell level in rich growth medium (***Dion et al., 2019***). We did not observe lysis in our growth conditions, possibly due to the reduced temperature and poorer growth medium, despite the drastic changes of cell width and mass density. Our finding demonstrates that crowding is not a limiting factor for growth in our growth conditions.

In conclusion, dry-mass density decreases with increasing cell width, both at the single-cell and at the population level, without affecting biomass growth rate.

### Inhibition of peptidoglycan insertion decouples surface growth from biomass growth

Next, we studied how surface growth is coupled to biomass growth mechanistically. Cell-wall insertion is generally thought to limit surface growth (***Kawai et al., 2009**; **Daniel and Errington, 2003**; **Rojas et al., 2017***). To test the potentially rate-limiting role of peptidoglycan synthesis for surface growth, we first inhibited cell-wall insertion by treating cells with the antibiotic vancomycin, which binds to the D-Ala-D-Ala terminus of peptidoglycan precursor molecules and thus inhibits cross-linking of new peptidoglycan material (***Barna and Williams, 1984***). To monitor single-cell growth we studied cells during time-lapse microscopy and added the drug to the pad about 30 min after placing cells on the microscope, similar to our nutrient-shift experiments (Figure 1 E,F).

Drug treatment leads to a sudden reduction of surface expansion about 10 min after adding the drug, while mass growth is affected much less (Figure 3A). The same behavior is observed at the single-cell level (Figure 3 - figure supplement 1A-B). The 10 min delay between drug addition and reduction of growth rates is at least partially due to the time it takes for the drug to diffuse through the agarose pad (see also next section). Accordingly, ***S**/**M*** decreases and biomass density increases in inverse proportion to ***S**/**M*** (Figure 3A, Figure 3 - figure supplement 1C-E). See Figure 3 - figure supplement 1A-B for single-cell traces.

**Figure 3.**
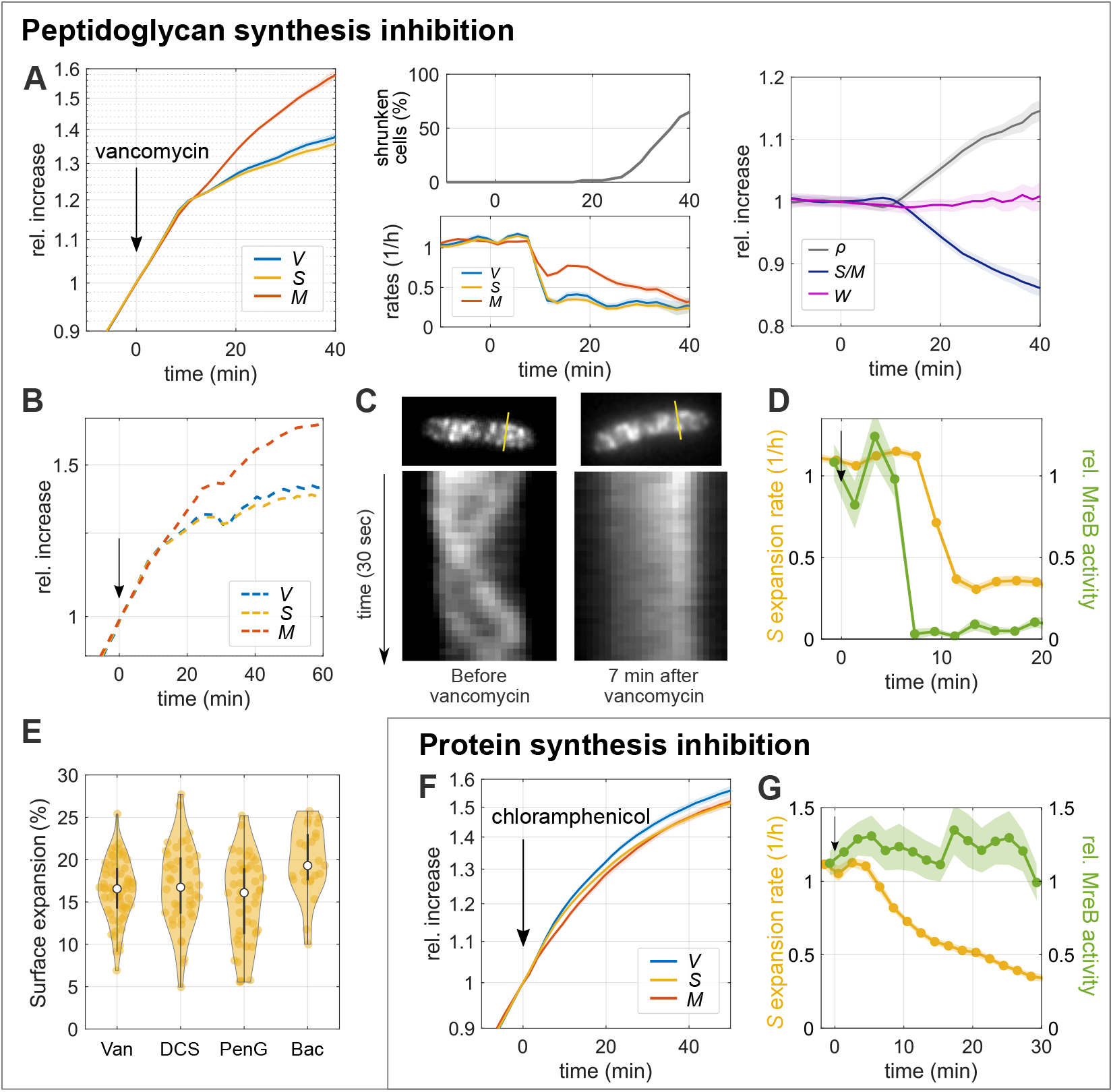
Inhibition of peptidoglycan insertion decouples surface growth from biomass growth. **A-B:** Single-cell time lapse of filamenting cells (bAB56) grown in S7_50_+GlcCaa medium and treated with vancomycin (50 μg/mL), which was added on top of the agarose pad at time = 0. To avoid cell division, MciZ was induced 30 min prior to microscopy. Relative increase (**left**) and rates (**middle-bottom**) of volume, surface, and dry mass. After 20 min, a fraction of cells starts to shrink in surface area (**middle-top**) and lose part of their mass (see also Figure 3 - figure supplement 2). **Right:** Relative change of dry-mass density, surface-to-mass ratio, and width. (Solid lines + shadings = average ± SE) **B:** Relative increase of volume, surface and dry mass for a representative single cell. **C:** Kymographs of MreB-GFP rotation in filamenting cells (bYS19) during 30 sec movie before and 7 min after vancomycin addition (as in A) along the lines indicated in MreB-GFP snapshots (top). **D:** Comparison of surface expansion rate (yellow; as in(A)) and relative MreB activity (total length of MreB tracks divided by projected cell area and movie duration; green) of vancomycin-treated cells to that of non-perturbed cells. (Solid lines + shadings = average ± SE) **E:** Residual surface expansion after MreB motion was arrested by Van (vancomycin 50μg/mL), DCS (D-cycloserine 10 mM). PenG (Penicillin G 0.5 mg/mL), Bac (Bacitracin 0.5 mg/mL). Experiments were performed in the same way as A and C (yellow dots = single-cell values; white circles = median; grey rectangles = interquartile range). **F:** Single-cell time lapse of filamenting cells (bAB56) treated with chloramphenicol (100μg/mL). Otherwise the same as in A. **G:** Comparison of surface expansion rate (yellow: as in (F)) and relative MreB activity (green) of chloramphenicol-treated cells to that of non-perturbed cells. (Solid lines + shadings = average ± SE)

About 10 min after the reduction of surface growth rate, some cells shrink in surface area and volume (Figure 3B), demonstrating a transient loss of envelope integrity and osmotic pressure. Consistently, many cells also lose a small fraction of their mass (Figure 3 - figure supplement 2). Interestingly, though, many cells continue to grow after such events (Figure 3 - figure supplement 2, Figure 3 - video 1).

We observed the same qualitative behavior when targeting cell-wall synthesis with different drugs that inhibit peptidoglycan-precursor synthesis (D-cycloserine), precursor transport (bacitracin), or cell-wall cross-linking (penicillin G) (Figure 3 - figure supplement 3). Thus, proper cell-wall insertion is apparently required for the maintenance of ***S**/**M*** during growth.

### Cell-envelope expansion can proceed in the absence of cell-wall insertion or protein expression

While cell-wall insertion is apparently required for the coordination between surface and biomass growth, cells still continue growing in surface area after drug treatment, even if at a reduced rate (Figure 3A, Figure 3 - figure supplement 3). This observation suggests that autolytic activity responsible for surface growth continues after cell-wall insertion is inhibited. To quantify the amount of residual surface growth after arrest of cell-wall insertion, we first aimed to identify the approximate time when cell-wall insertion is arrested. We therefore tracked the movement of MreB-actin filaments, using the mutant strain (bYS19) that expresses a GFP-MreB protein fusion from the native locus (***Dion et al., 2019***). MreB rotation depends on cell-wall insertion (***Garner et al., 2011**; **Domínguez-Escobar et al., 2011***), and the number of moving MreB filaments is linearly correlated with the rate of cell-envelope growth, if growth rate is modulated through nutrient quality (***Sun and Garner, 2020***). We thus used MreB rotation as a readout for ongoing cell-wall insertion. More specifically, we measured the product of MreB-filament density (number of fllaments per cell-contour area) times average speed, by simply summing up all MreB-track lengths and dividing by 2D cell area (contour area) and movie duration. We refer to this quantity as ‘MreB activity’. However, since diffraction-limited microscopy impedes the detection of all MreB filaments, we restricted our interpretation to large relative changes of MreB activity.

In agreement with previous observations (***Garner et al., 2011**; **Domínguez-Escobar et al., 2011***), all drugs used here (vancomycin, penicillin G, D-cycloserine, and bacitracin) stop MreB motion within 4-8 min after adding the drug on top of the agarose pad (Figure 3C, D, Figure 3 - figure supplement 4, Figure 3 - video 2, Figure 3 - video 3). The delay is likely entirely due to the time it takes the drugs to diffuse to the cells. Our experiments suggest that cell-wall insertion is either inhibited or drastically reduced at the time of MreB-motion arrest, while cell-wall expansion continues by about 10 - 20% during the residual time of observation (Figure 3E).

By comparing the time-dependent rates of surface growth and MreB activity at early times after different drug treatments, we also found that surface expansion proceeds at an unperturbed rate for about 2-6 min after MreB motion has stalled (Figure 3D, Figure 3 - figure supplement 4), in qualitative agreement with previous observations (***Misra et al., 2013***) of the experimental data by ***Garner et al.** (**2011***). Thus, cell-surface growth and cell-wall insertion are not strictly coupled.

Next, we wondered whether we could find additional conditions under which cell-wall expansion and cell-wall insertion are decoupled. It was previously shown at the population-level that the inhibition of protein translation through chloramphenicol leads to a rapid reduction of biomass growth (based on turbidity), while peptidoglycan synthesis continues (***Chung, 1967***) and cell-wall thickness increases (***Miller et al., 1967***). Here, we investigated single cells treated with 100 μg/mL of chloramphenicol, which completely inhibits protein translation (Figure 3 - figure supplement 5), in time-lapse microscopy. In agreement with ***Chung** (**1967***) we observed that mass growth and cell-surface growth are strongly reduced (Figure 3F, Figure 3 - figure supplement 5, Figure 3 - video 4), while MreB activity remains high (Figure 3G, Figure 3 - video 5). Interestingly, cell-envelope and biomass growth remain coupled despite the severe perturbation. This coupling is also observed if we correct our calculation of cytoplasmic surface area and mass for cell-wall thickening (Figure 3 - figure supplement 6), a consequence of continued cell-wall insertion. Thus, our observations suggest that cells can regulate surface expansion through a pathway that is different from cell-wall insertion. Furthermore, our observation also demonstrates that the insertion of new envelope proteins is not required and therefore not rate-limiting for surface growth.

Together, the rates of cell-wall expansion and cell-wall insertion are not strictly coupled, suggesting that the activity of cell-wall-cleaving hydrolases is controlled by a pathway that is independent of cell-wall insertion.

### Inhibition of membrane synthesis also decouples surface expansion and mass growth

A different envelope component was recently demonstrated to have an important influence on cell-envelope growth (***Rojas et al., 2017***): the cytoplasmic membrane. We therefore investigated how a modulation of membrane synthesis affects surface expansion. First, we treated cells with cerulenin, which inhibits fatty-acid synthesis (***Omura, 1976***) and phospholipid insertion in the closely related *Bacillus amyloliquefaciens (**PATON et al., 1980***) and also in *E. coli*, at about 100 *μ*g/ml.

As for cell-wall-synthesis inhibitors, we first added cerulenin at about 100 *μ*g/ml to the top of an agarose pad, which then reaches the cells through diffusion on the timescale of few minutes. Within 7 min after drug addition, both surface expansion and mass growth are reduced (Figure 4A, Figure 4 - video 1), qualitatively similar to the inhibition of peptidoglycan insertion (Figure 3A). However, different from the inhibition of cell-wall insertion, cerulenin does not cause visible lysis or partial loss of mass and turgor. Since surface growth is affected more severely than mass growth, ***S**/**M*** decreases and mass density increases. We observed a very similar behavior when the drug was already contained in the agarose pad, that is, when the cells were immediately exposed to the drug at its final concentration (Figure 4 - figure supplement 1), apart from the initial diffusion-caused delay (Figure 4A). Thus, proper membrane synthesis is required for the maintenance of ***S**/**M***.

**Figure 4.**
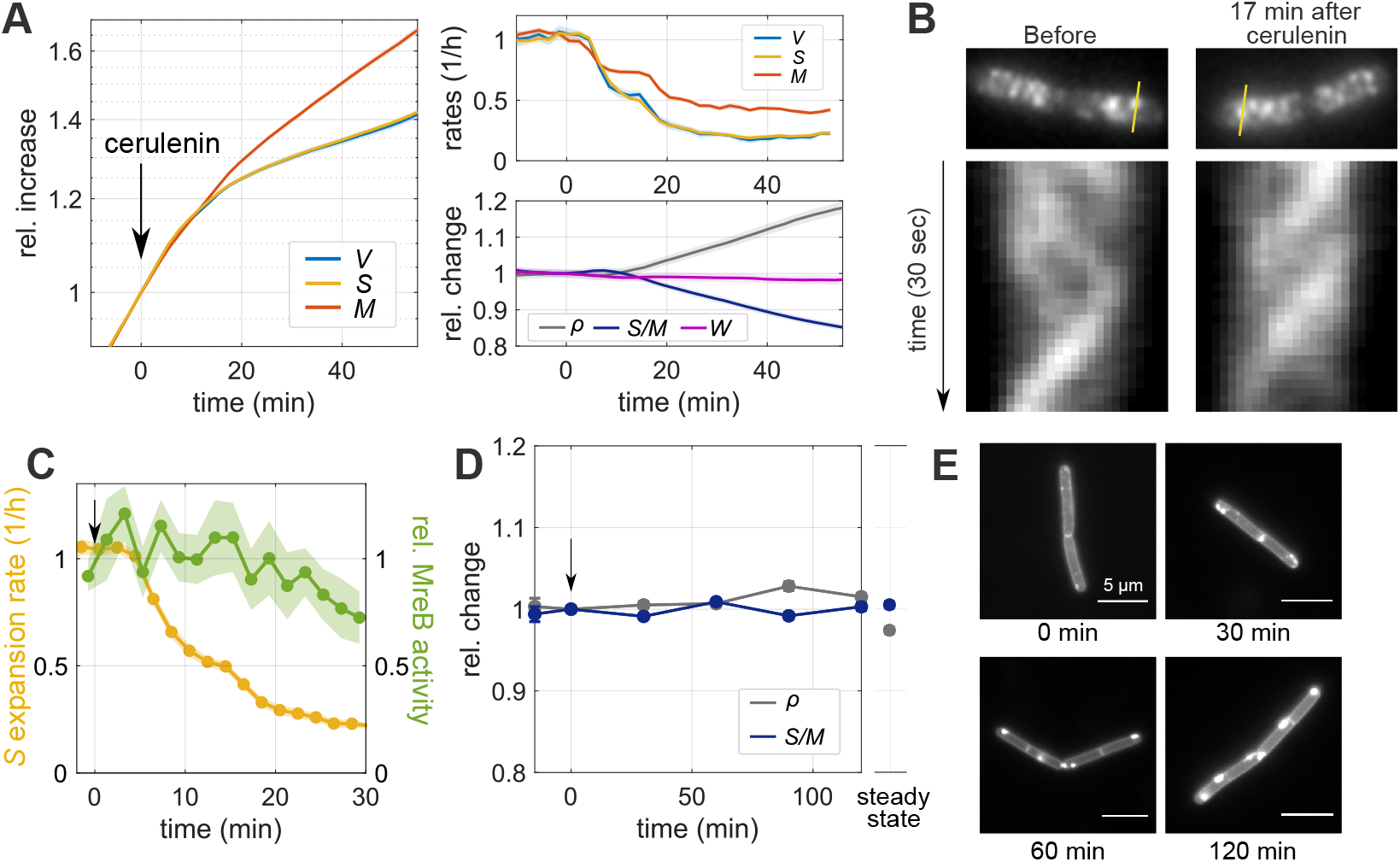
Inhibition of membrane synthesis decouples surface expansion and mass growth independently of peptidoglycan synthesis. **A:** Single-cell time lapse of filamenting cells (bAB56) grown in S7_50_+GlcCaa and treated with cerulenin 100μg/mL (added to the agarose pad at time = 0). Relative increase (left) and rates (top right) of volume, surface area, and dry mass. Relative change (bottom right) of dry-mass density, surface-to-mass ratio, and width. (Solid lines + shadings = average ± SE) **B:** Kymographs of MreB-GFP rotation in filamenting cells (bYS19) during 30 sec movie before and 17 min after cerulenin addition along the lines indicated in MreB-GFP snapshots (top). **C:** Comparison of surface expansion rate (yellow; as in(A)) and relative MreB activity (green) of cerulenin-treated cells to that of non-perturbed cells, similar to Figure 3D. (Solid lines + shadings = average ± SE) **D:** Relative change of average dry-mass density and surface-to-mass ratio upon overexpression of AccDA (bSW305: *amyE*::pXyl-*accDA*) by addition of xylose (10 mM) at time = 0 in LB medium (average ± SE). Every point represents the average obtained from snapshots of batch-culture-grown cells. **E:** AccDA overexpression in bSW305 (the same experiment in D) leads to the accumulation of excess membrane according to staining with the membrane dye MitoTracker green.

The increase of mass density after cerulenin treatment is consistent with previous work by ***Mindich** (**1970***), who demonstrated that buoyant mass density of cells inhibited in membrane synthesis by glycerol starvation of a glycerol auxotroph is visibly increased after about one generation time.

Given the similarity between cerulenin treatment and our inhibition of cell-wall synthesis (Figure 3A), we wondered whether cerulenin might lead to a reduction of surface growth by affecting cell-wall insertion. We thus monitored MreB-GFP activity as in Figure 3C-D. We found that MreB activity is not affected for at least 15 min after the initial reduction of surface expansion (Figure 4B,C, Figure 4 - video 2), followed by a mild reduction during the remaining observation time. We confirmed these results with a complementary method developed by some of us (***Dion et al., 2019***). The method yields area density and speed of moving MreB filaments based on total-internal reflection microscopy (TIRF) and a subsequent kymograph-based analysis (Figure 4 - figure supplement 2). The method, which was previously shown to compare well with independent high-resolution structured-illumination microscopy (***Dion et al., 2019***), confirms our results.

Our observations therefore suggest that membrane synthesis affects surface expansion independently of peptidoglycan insertion. Our finding is in agreement with previous work from ***Mindich** (**1970**)*: He showed that cell-wall synthesis continues after membrane synthesis is inhibited by glycerol starvation, according to the incorporation of radioactive alanine (***Mindich, 1970***).

Similar to chloramphenicol treatment (Figure 3F-G), the cell wall likely thickens during cerulenin treatment. Based on the small effect in chloramphenicol-treated cells on the coupling of surface and mass (Figure 3 - figure supplement 6), cell-wall thickening should also not affect the decoupling of surface and mass in cerulenin-treated cells.

The rate of mass growth slows down as time progresses. We initially speculated that this decrease might be a consequence of increased crowding. However, when inspecting the relationship between mass growth rate and mass density at the single-cell level, we observed no visible correlations (Figure 4 - figure supplement 3), supporting our conclusion drawn from the constant growth rate after modulation of aPBP levels (Figure 2F): Mass density is likely not rate-limiting for mass-growth rate.

Next, we studied a previously described mutant (bSW305, *amyE*::pXyl-*accDA*) that overproduces membrane lipids when grown in LB medium (***Mercier et al., 2013***). If the cytoplasmic membrane was rate-limiting for surface growth, we would expect an increase of ***S**/**M*** upon *accDA* induction. However, we did not observe any change of ***S**/**M*** both during steady-state growth or during the first 2 h after *accDA* induction (Figure 4D), for a control see Figure 4 - figure supplement 4). At the same time, we observed apparent excess cytoplasmic membrane according to membrane staining with MitoTracker green (Figure 4E) demonstrating increased membrane production in agreement with previous investigations by electron microscopy (***Mercier et al., 2013***). Thus, while proper membrane physiology is apparently important for cell-envelope growth, independently of peptidoglycan insertion, excess membrane production does not lead to an increase of surface growth.

### Cells actively restore the surface-to-mass ratio through transiently increased surface growth

Since cells robustly maintain ***S**/**M*** during steady-state growth and during nutrient shifts (Figure 1), we wondered whether and how they restore ***S**/**M*** after perturbation. To perturb ***S**/**M***, we treated wild-type cells with cerulenin in batch culture for 30 min, which is expected to lead to a decrease of ***S**/**M*** by about 10% (Figure 4A). We then washed out the drug and studied cells during a time-lapse movie on an agarose pad (Figure 5A, Figure 5 - figure supplement 1, Figure 5 - video 1). During the first 20 min, ***S**/**M*** continues to decrease while mass density increases, possibly because cerulenin binds its target (*β*-ketoacyl-ACP synthase) irreversibly (***D’agnony et al., 1973***). However, during the subsequent 60 min, which corresponds to one mass doubling, ***S**/**M*** returns to its steady-state (pre-treatment) value, and mass density decreases accordingly. Remarkably, ***S**/**M*** changes almost deterministically at the single-cell level, both during treatment and recovery (Figure 5 - figure supplement 2). Thus, ***S**/**M*** is likely restored even at the single-cell level.

**Figure 5.**
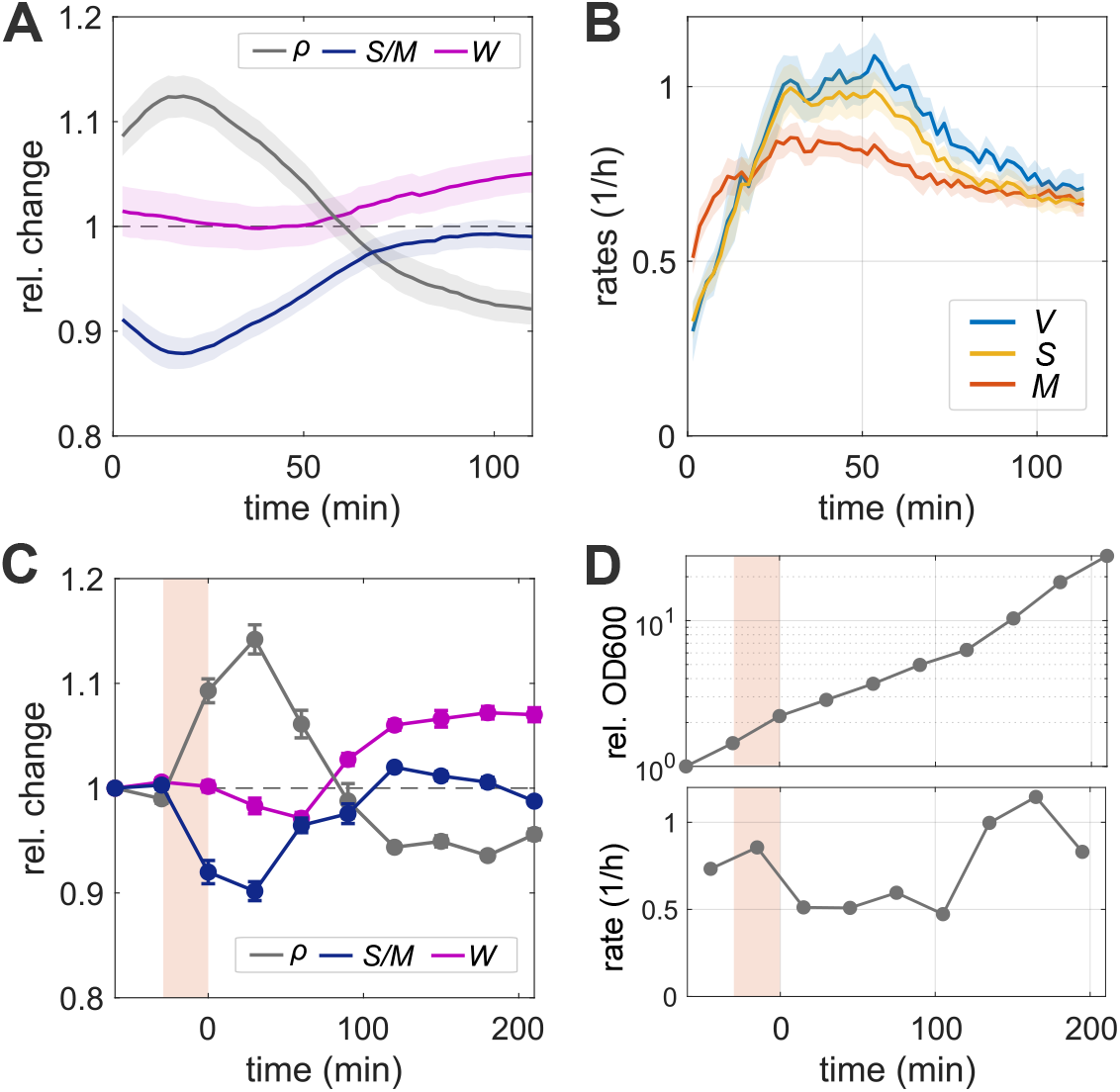
Cells restore the surface-to-mass ratio through transiently increased surface growth. **A-B:** Single-cell time lapse of bAB56 cells during recovery from 30 min cerulenin treatment (100 μg/ml) on an agarose pad in S7_50_+GlcCaa. **A**. To avoid cell division, MciZ was induced 50 min prior to microscopy. Relative change of dry-mass density, surface-to-mass ratio, and width, normalized with respect to steady-state conditions. **B**. Rates of volume, surface, and dry mass growth (Solid lines + shadings = average ± SE). **C-D:** The same recovery experiment as in (A-B) in batch culture, followed before, during, and after cerulenin treatment. The culture was back diluted to keep optical density < 0.3. **C**. Relative changes of average dry-mass density, surface-to-mass ratio, and width (average ± SE) obtained from single-cell snapshots as a function of time (shaded region: duration of cerulenin treatment; time =0 corresponds to the time of washout). **D**. Relative change of optical density (OD600), after normalization for backdilution, and growth rate.

Consistently with the recovery of ***S**/**M***, we observed that surface growth rate increases continuously from the reduced level observed during treatment in Figure 4A, to a level that transiently exceeds mass growth by about 20% (Figure 5B), before returning to the rate of mass growth. Notably, mass growth rate increases rapidly within the first 10 min, but then remains lower than the steady-state growth rate, which is about 1/h (Figure 1D). Our observations are consistent with the previous study of ***Mindich** (**1970***), which shows rapid resumption of membrane and biomass synthesis during re-feeding of glycerol after transient glycerol starvation.

We observed the same behavior in snapshots of cells growing in batch culture (Figure 5C-D): ***S**/**M*** drops by about 10% during the 30 min cerulenin treatment, while mass density increases by the same relative amount. After washout, ***S**/**M*** first continues to decrease and then returns to its steady-state value within one mass-doubling time according to optical density (Figure 5C-D), in agreement with the time-lapse experiment (Figure 5A-B). Mass growth rate recovers to the pre-treatment value after about 120-150 min.

The time-dependent recovery of ***S**/**M*** is qualitatively similar to what we expect from the empirical surface growth law that relates d***S***/d*t* linearly to d***M***/d*t* (***Oldewurtel et al., 2019***). According to this model, ***S**/**M*** should return asymptotically to its pre-shift value, with a rate equal to the instantaneous mass growth rate (Figure 5B). During the period of ***S**/**M***-recovery, the expected timescale of recovery is thus about [dlog(***M***)/d*t*]^-1^ ≈ 75 min (the time to recover 63% of the transient loss of ***S**/**M***). Here, we find that ***S**/**M*** recovers even faster, with an approximate recovery time of 45 min. This speedup might be due to the accumulation of other cell-envelope material during cerulenin treatment, which is reminiscent of previous reports of ‘supergrowth’ in the yeast *Schizosaccharomyces pombe* after transient reduction of the surface growth (***Knapp et al., 2019***). However, more work will be required to understand this phenomenon quantitatively.

The reduction of mass density by about 5% after about 100 min in both time-lapse and batch experiments can be reconciled with a similar systematic increase in average cell width that persists for longer than our observation time. While we don’t know the cause of this increase, this observation demonstrates that ***S**/**M*** robustly returns to its pre-treatment value despite a systematic reduction of mass density.

## Discussion

In conclusion, the Gram-positive bacterium *B. subtilis* controls cell-volume growth indirectly, by increasing surface area in proportion to biomass growth, qualitatively in the same way as the Gramnegative *E. coli (**Oldewurtel et al., 2019***). More specifically the surface-to-mass ratio ***S**/**M*** remains almost constant, independently of cell-to-cell variations of cell width or instantaneous growth rate. Since average width of *B. subtilis* does not systematically change in different nutrient conditions (***Sharpe et al., 1998***), surface-to-mass coupling can guarantee density homeostasis during growth. However, the constancy of ***S**/**M*** is broken if either cell-wall insertion or membrane synthesis is perturbed. Thus, both of these processes are required for proper surface-to-mass coupling and therefore for volume regulation. Once the inhibition of surface growth is relieved, cells rapidly recover their steady-state surface-to-mass ratio by growing faster in surface than in mass.

While ***S**/**M*** is independent of stochastic cell-to-cell variations of width, a systematic increase of average width by PonA overexpression leads to a reduction of the average value of 〈***S**/**M***〉 and a decrease of width through deletion of PonA or all class-A PBPs leads to an increase of 〈***S**/**M***〉 (Figure 2D). Different mechanisms might be responsible for this correlation: First, the modified ratio of MreB-based and class-A PBP-based cell-wall insertion is known to affect cell-wall architecture (***Dion et al., 2019**; **Turner et al., 2013***), which might then affect autolytic activity. Second, changes of cell width and dry-mass density likely affect mechanical envelope stresses, which might also affect autolytic activity (***Koch, 1983***). PonA expression or depletion is also known to affect the expression of different cell-wall-related proteins through the sigma factor σ^|^ (***Patel et al., 2020***). Finally, the change of 〈***S**/**M***〉 could also be the result of a yet unknown feedback between mass density and cell-wall expansion. Further work will be required to understand the effect of class-A PBP expression on surface growth.

The conserved indirect control of cell-volume growth through surface-to-mass coupling across Gram-negative and Gram-positive bacteria is remarkable given their fundamentally different envelope architectures. We previously reasoned, based on our findings in *E. coli **(Oldewurtel et al., 2019***), that the coupling of surface and mass might have a metabolic origin: Constancy of ***S**/**M*** would come about if cells devoted a constant fraction of newly acquired mass to one or multiple envelope components whose production are rate-limiting for surface growth. Here, we found that both cell-wall insertion and membrane synthesis are required for the maintenance of ***S**/**M*** in *B. subtilis*, thus providing two links between metabolism and cell-envelope expansion.

The dependency of ***S**/**M*** on cell-wall insertion is qualitatively different from *E. coli*, which expands surface area independently of cell-wall insertion (***Oldewurtel et al., 2019***). However, when perturbing cell-wall insertion in *B. subtilis*, we also observed significant deviations between cell-wall expansion and MreB-based cell-wall insertion: Most notably, inhibiting peptidoglycan insertion causes a reduction of MreB rotation only after a short but significant delay of 2-6 min (Figure 3D, Figure 3 - figure supplement 4), and cell-wall expansion continues at a reduced rate even after MreB rotation is completely arrested. Furthermore, cells start to shrink and lose part of their mass or even lyse after about 20 min after drug exposure. These findings suggest that autolytic enzymes, which are physically responsible for cell-wall growth, do not directly depend on cell-wall insertion but are controlled through an unknown signal, which, in turn, is affected by peptidoglycan insertion.

Inhibition of fatty-acid biosynthesis through cerulenin treatment leads to an equally rapid reduction of surface growth as the inhibition of cell-wall insertion (Figure 4A). The cytoplasmic membrane is thus arguably equally important for the regulation of surface growth and thus for cell-volume regulation as the cell wall. Previously, it was demonstrated by different groups that membrane tension and membrane fluidity are important factors that modulate cell-wall insertion, which might affect surface growth (***Müller et al., 2016**; **Zielińska et al., 2020**; **Rojas et al., 2017***). Here, we found that the inhibition of fatty-acid synthesis reduces surface growth through a mechanism that is different from cell-wall insertion (Figure 4A-C), in qualitative agreement with (***Mindich, 1970***). However, while membrane insertion is apparently required for proper surface growth, visible overproduction of membrane upon overexpression of the Acetyl-CoA carboxylase components AccDA does not lead to an increase rate of surface growth (Figure 4D,E), even if excess membrane likely contributes to more surface area after protoplast formation (***Mercier et al., 2013***). This finding suggests that the flux of total membrane lipids is also not the sole rate limiting envelope component. However, it remains possible that the synthesis of specific lipids has a rate-limiting role for envelope growth. The role of the cytoplasmic membrane thus deserves further investigation in the future.

How membrane synthesis and cell-wall synthesis are linked to biomass growth is a question that remains fundamentally not understood in any bacterium. The first committed steps of fatty-acid and phospholipid synthesis are likely the major pathway elements for the control of membrane synthesis (***Cronan Jr and Rock, 2008**; **Dowhan, 2013**; **Noga et al., 2020**; **Rock and Cronan, 1996**; **Cronan Jr and Waldrop, 2002***). However, the signals responsible for controlling their activities largely remain to be identified (***Noga et al., 2020***). Furthermore, while multiple gene-regulatory feedbacks for cell-wall metabolism have been identified (***Patel et al., 2020***), the question of how cell-wall metabolism and mass growth are robustly coupled remains open. Recent work from some of us (***Sun and Garner, 2020***) has identified PrkC as an important regulator of MreB-based cell-wall insertion. ***Sun and Garner** (**2020***) suggested that PrkC senses the availability of lipid II, the precursor of peptidoglycan synthesis, and thus regulates the number of moving MreB filaments. However, it remains to be investigated if and how lipid II levels are an important factor to coordinate cell-wall insertion and biomass growth.

Interestingly, inhibition of cell-wall insertion or fatty-acid synthesis does not only reduce the rate of surface growth but also affects biomass growth rate (Figure 3A, Figure 4A). Mass-growth rate drops synchronously with the reduction of surface growth (at our time resolution of 2 min), even if the reduction is less pronounced. At this time point, mass density is not visibly affected. The reduction of mass growth rate is thus not a response to increased crowding but likely triggered by an active signaling pathway or possibly because of deficiencies in nutrient uptake as speculated by ***Mindich** (**1975***).

Previous work demonstrates that the stringent response is required for cell survival after ceru-lenin treatment in both *B. subtilis (**Pulschen et al., 2017***) and *E. coli (**Vadia et al., 2017***), suggesting a potential role in reducing biomass growth in response to the arrest of membrane synthesis. However, on the generation time scale, the stringent response is not required for the reduction of biomass growth in *B. subtilis (**Pulschen et al., 2017***). Thus, one or multiple different pathways must be responsible. ***Sun and Garner** (**2020***) proposed that levels of the peptidoglycan precursor lipid II affect both cell-wall insertion and mass growth through the kinase PrkC. Interestingly though, we found that the reduction of mass growth coincides with the reduction of surface growth and not with the time of MreB-rotation arrest. There are thus likely additional links between surface growth and mass growth. In any case, surface-to-mass coupling appears to be bidirectional, with biomass growth affecting surface growth, but surface growth also affecting biomass growth, even if to a lesser extent.

At long times after the inhibition of cell-wall insertion or membrane biosynthesis, mass density increases due to the differences between mass and surface growth rates. We initially speculated that increased crowding might cause a decrease of mass growth rate observed during late times of drug treatment. However, to our surprise, we found that mass growth rate remained constant during steady-state exponential growth at different levels of PonA induction or class-A PBP deletion (Figure 2F), which can cause similarly high levels of dry-mass density as vancomycin/cerulenin treatment. Furthermore, single-cell mass growth rate does not visibly correlate with mass density at different times after cerulenin treatment (Figure 4 - figure supplement 3). Mass density and crowding are considered important determinants of biomass growth rate, for example through their effect on the diffusion of tRNA complexes (***Klumpp et al., 2013***) or through a potential effect on the density of metabolites (***Vazquez, 2010**; **Dourado and Lercher, 2020***). However, constancy of growth rate despite strong differences in density suggests that crowding or density are not limiting factors for growth rate in *B. subtilis* in our growth conditions.

## Materials and Methods

### Key Resources Table

| Reagent type (species) or resource | Designation | Source or reference | Identifiers | Additional information |
| --- | --- | --- | --- | --- |
| strain, (*B. subtilis*) | Wildtype |  | PY79 |  |
| strain, (*B. subtilis*) | bAB56 | This work | *mciZ*::spec pHyperSpank-*mciZ* |  |
| strain, strain background (*B. subtilis*) | bMD586 | (Dion et al., 2019) | *yhdG*::cat pHyperSpank-*ponA*, *ponA*::kan (Δ*ponA*) | |
| strain, (*B. subtilis*) | bKY42 | This work | *ponA*::kan |  |
| strain, (*B. subtilis*) | bSW164 | This work | *pbpD*::lox72, *pbpG*::lox72, *pbpF*::lox72, *ponA*::kan, *amyE*::spec pSpac-*mciZ* |  |
| strain, (*B. subtilis*) | bYS19 | (Dion et al., 2019) | *mreB*::*mreB*-msfGFPsw |  |
| strain, (*B. subtilis*) | bSW305 | This work | *amyE*::tet pXyl-*accDA* |  |
| software | MATLAB | The MathWorks, Inc. |  |  |
| software, algorithm | Morphometrics | (Ursell et al., 2017) |  |  |
| software, algorithm | BlurLab | (Ursell et al., 2017) |  |  |
| software | Fiji | (Schindelin et al., 2012) |  |  |
| software, algorithm | Trackmate | (Tinevez et al., 2017) |  |  |
| software, algorithm | MicroManager | (Edelstein et al., 2010) |  |  |

### Growth conditions and sample preparation

Cell cultures were grown from a single colony in liquid media at 30°C in a shaking incubator. We used three different growth media: LB (Luria-Bertani Miller medium), S7_50_+Glc (minimal medium as described in (***Jaacks et al., 1989***), except that 0.4% glucose and 20 mM glutamate were used rather than 1% and 0.1%, respectively), and S7_50_+GlcCaa (S7_50_+Glc supplemented with 0.4% casamino acids). Before microscopy, we kept cultures in exponential phase for >10 mass doublings at OD600 < 0.3 through back-dilution.

For single-cell snapshots or time-lapse movies we immobilized cells under a pre-warmed agarose pad (1.5% UltraPure Agarose (16500-500, Invitrogen)). Microscopy in a 30°C incubator was started within 3 min after cells were placed on the agarose pad.

For time-lapse movies, images were taken every 2 min if not specified. To avoid cell division, we started inducing *mciZ* from an IPTG-inducible promoter by adding 1 mM IPTG to the culture prior to microscopy and we added 1 mM IPTG in the agarose pad during microscopy (For the time of IPTG addition, see Supplementary File 1). While MciZ inhibits the formation of new septa, about 1/3 of cells still contained non-complete septa when placing cells on agarose pads, according to FM 4-64-based membrane staining. Thus, some of the cells analyzed are likely separated by a septum at the end of the time lapses, even if they are not visibly separated according to their contour. For simplicity we considered possibly chained cells as single cells.

To stain the cytoplasmic membrane, 1 μg/mL of FM 4-64 Dye (ThermoFisher, T13320) was contained in agarose media. For perturbations, we added the following compounds to the top of the agarose pad: alpha methylglucoside (0.4%), vancomycin (50 μg/mL), D-cycloserine (10 mM), penicillin G (500 μg/mL), bacitracin (500 μg/mL), chloramphenicol (100 μg/mL), cerulenin (100 μg/mL). Biological replicates result from independent cultures starting from separate colonies.

### Strain construction

All strains used in this study derive from the wildtype PY79. Strains, plasmids, DNA fragments and oligonucleotides are all described in Supplementary File 2.

#### bAB56

(*mciZ*::spec-pHyperSpank-*mciZ*) was generated upon transformation of PY79 with a four-piece Gibson assembly reaction that contained the following amplified fragments: upstream of the *mciZ* gene; spectinomycin-resistance cassette loxP-spec-loxP; the *lacI* gene and the pHyperSpank promoter with an optimized ribosomal binding sequence; the *mciZ* coding region and downstream sequence.

#### bKY42

(*ponA*::kan) was generated upon transformation of PY79 with genomic DNA from bMD586 (***Dion et al., 2019***).

#### bSW164

(*pbpD*::lox72, *pbpG*::lox72, *pbpF*::lox72, *ponA*::kan, *amyE*::spec-pSpac-*mciZ*) was generated upon successive rounds of transformation of PY79 with genomic DNA from strains bMK258 (*pbpD*::erm), bMK260 (*pbpG*::erm), bMK270 (*pbpF*::erm), described below, and bSW99 (*amyE*::spec-pSpac-*mciZ*) (***Hussain et al., 2018***) and bMD599 (*ponA*::kan) (***Dion et al., 2019***). After each transformation, the erythromycin-resistance cassette was removed with plasmid pDR244 (***Koo et al., 2017***). bMK258 (*pbpD*::erm), bMK260 (*pbpG*::erm), and bMK270 (*pbpF*::erm) were generated upon transformation of PY79 with a three-piece Gibson assembly reaction that contained the following amplified fragments: upstream of the respective gene to be deleted; erythromycin-resistance cassette loxP-erm-loxP; downstream of the respective gene.

#### bSW305

(*amyE*::tet-pXyl-*accDA*) was generated upon transformation of PY79 with a five-piece Gibson assembly reaction that contained the following amplified fragments: upstream of the *amyE* gene; tetracyclin-resistance cassette loxP-tet-loxP; the xylR gene and the pXylA promoter with an optimized ribosomal binding site; the *accDA* coding region; downstream of the *amyE* gene.

### Microscopy

Except for the TIRF-based MreB density measurements (Figure 4 - figure supplement 2), microscopy was carried out on a Nikon Ti-E inverted phase-contrast and epi-fluorescence microscope that is additionally equipped with a module for spatial light interference microscopy (SLIM) (***Wang et al., 2011***) as described in detail in (***Oldewurtel et al., 2019***). The microscope is equipped with a temperature chamber (Stage Top incubator, Okolab) set to 30°C, a Nikon Plan Apo 100x NA 1.45 Ph3 Objective, a solid-state light source (Spectra X, Lumencor Inc. Beaverton, OR), a multiband dichroic (69002bs, Chroma Technology Corp., Bellows Falls, VT), and with excitation (485/25, 560/32) and emission (535/50, 632/60) filters for GFP and FM 4-64 imaging, respectively. Epi-fluorescent images were acquired with a sCMOS camera (Orca Flash 4.0, Hamamatsu) with an effective pixel size of 65 nm, while phase-contrast and quantitative phase images were obtained with another CMOS camera (DCC3260M, Thorlabs) with an effective pixel size of 87 nm. For SLIM measurements we took six consecutive images with a phase delay of nπ/2, where n=[1,2,3,4,5,6], with 200 ms exposure each. Out of these, we obtained three phase images (from images 1-4, 2-5, and 3-6, respectively), and took the average to obtain the final phase image. Including delays due to software, the acquisition of one final phase image took <3 seconds. Micro-manager was used to control the microscope and acquire images within MATLAB.

For the TIRF-based investigation of MreB rotation shown in Figure 4 - figure supplement 2, which was carried out in Ethan Garner’s lab, microscopy was carried out on a Nikon Ti phase-contrast and TIRF microscope, equipped with temperature control, a Nikon 100X NA 1.45 objective, and a sCMOS camera (Orca Flash 4.0, Hamamatsu) with an effective pixel size of 65 nm. Nikon NI Elements was used to control the microscope.

### Measurement of cytoplasmic contour, dimensions, surface area, and volume

Cell dimensions were obtained from phase-contrast images acquired using the SLIM module, essentially as described previously (***Oldewurtel et al., 2019***). Specifically, we used the MATLAB-based tool Morphometrics (***Ursell et al., 2017***) to determine cell contours. The image-formation process through the microscope, but also the contour-finding routines of Morphometrics can bias and distort the contour. We correct and calibrate for this based on epi-fluorescence images of cells stained with the fluorescent membrane stain FM 4-64. Since the calibartion is generally cell-shape dependent, we collected FM 4-64 images for wild-type cells and ponA-expressing mutant cells (*yhdG*::cat-pHyperSpank-*ponA*,*ponA*::*kan*) with different levels of inducer grown in S7_50_+GlcCaa medium. For FM 4-64 image acquisition, we focused on the middle of the cell based on phase-contrast microscopy through the epi-fluorescence port, which yields a sharper cell contour than the SLIM module. To correct these images for diffraction, we simulated membrane-stained cells as described (***Oldewurtel et al., 2019***) using the MATLAB based tool BlurLab (***Ursell et al., 2017***) and using the point-spread function (PSF) of the microscope (based on 100 nm fluorescent beads (TetraSpeck, Thermo Fisher)). We applied the correction found in silico onto membrane contours obtained by Morphometrics to obtain the true (physical) contour of the periphery of the cytoplasm. In addition to the epi-fluorescence images, we obtained phase-contrast images of the same cells using the SLIM module. We then overlaid the measured contour of the phase-contrast cell with the corrected membrane contour obtained from the membrane dye and measured their respective offset as a function of cell width and as a function of the distance from the cell pole (Figure 1 - figure supplement 1B).

This correction was used to correct the contours of all cells measured with the SLIM module. Finally, given the calibrated contours of the cell, we used Morphometrics to apply a mesh-grid of 1 px (87 nm) step-size. This routine also gives the centerline of the cell, which is used to determine cell length. We then assume cylindrical symmetry around the centerline and infer cell surface and cell volume from the sum of the surfaces and volumes of truncated conical wedges with height and width given by the meshes.

For the confirmation of continued MreB activity after cerulenin treatment through TIRF microscopy (Figure 4 - figure supplement 2) we also segmented phase-contrast images using the Morphometrics tool. However, those data were not calibrated against FM4-64 images. This is not relevant for the calculation of instantaneous surface-growth rate or density of moving MreB filaments.

### Experimental quantification of cell-wall dry mass

Cultures of wildtype and bMD586 were grown to exponential phase in 1 liter of S7_50_+GlcCaa medium. For the bMD586 culture, *ponA* expression was induced by 1 mM IPTG. Once OD600 reached 0.3, cells were harvested and washed with Milli-Q water. The suspension of the cells in Milli-Q water was evenly divided into two. The harvested cells from one suspension were subjected to vacuum drying overnight and the dry weight was measured. The other suspension was subjected to sonication to break cells (complete cell disruption was confirmed by microscopy), and the insoluble fraction, which consist predominantly of cell wall, was washed by Milli-Q water. After vacuum drying overnight, the dry weight of the cell-wall fraction was measured. The cell-wall content (*ζ*) is calculated as dry weight of the cell-wall fraction per dry weight of total cell suspension.

### Calculation of cytoplasmic dry mass from quantitative-phase images

The cytoplasmic dry mass is calculated as

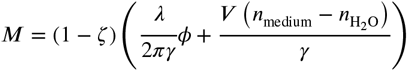

Here, *λ* = 635 nm is the central wavelength of light, *n*_medium_ and *n*_H_2_O_ are refractive index of the medium and water respectively, and *ζ ≈* 0.14 is the fraction of biomass occupied by the cell wall (obtained from bulk experiments; see previous section). For a correction of this value due to cell-wall thickening during treatment with either chloramphenicol (Figure 3F-G) or cerulenin (Figure 4A) see the subsequent section.

In case of experiments using an agarose pad we added *n*_agarose_ (= 0.0020) to *n*_medium_. We measured *n*_medium_ using a refractometer (Brix/RI-Chek, Reichert). *γ* is the refraction increment of the cell, estimated below, and *φ* is the integrated phase obtained from the phase image, detailed below. We defined cytoplasmic dry mass as all the dry mass other than cell wall fraction which also includes periplasmic molecules. Although our approach thus overestimates cytoplasmic dry mass by up to 5% when considering reported amount of periplasmic proteins (***Merchante et al., 1995***) and lipoteichoic acid (***Huff, 1982***), this does not affect relative change.

To calculate the refraction increment, we considered the reported composition of dry mass (***Bishop et al., 1967***) and reported values for refraction increments (***Theisen, 2000**; **Marquis, 1973**; **Barer and Joseph, 1954**; **Barer, 1956***) (Supplementary File 3). Within the uncertainty of the refraction increments for individual cell constituents, the weighted average refraction increment is between 0.175 - 0.182 mL/g. To account for the higher illumination wavelength of 635 nm used in our experiments, we further decrease the refraction increment by 1% (***Perlmann and Longsworth, 1948***). Thus, we arrive at the average refraction increment of *γ* = 0.177 mL/g, which we used for all conversions.

*φ* is the integrated phase obtained from the phase image, after correction for attenuation by optical artifacts of the microscope, notably the halo effect, as described previously (***Oldewurtel et al., 2019***). In brief, the integrated phase is underestimated by about two-fold, but the precise attenuation depends on cell geometry. To correct for this attenuation, we conducted computational simulations of phase images for every cell and every time point that are informed by the properties of the microscope and by the cytoplasmic contour. We then integrated the measured phase in simulated images and compared this value to the expected integrated phase from the simulation parameters (ground truth). This comparison yields an attenuation factor used to correct the underestimated integrated phase from experiments. We repeated this procedure for every cell and everytime point.

Strictly speaking, the attenuation factor should be calculated based on the the contour of the cell (rather than the contour of the cytoplasm). However, the attenuation factor changes by less than 1.5% if we assume a contour that is larger (in radial direction) by 56 nm, the sum of a potential periplasm (22 nm according to ***Matias and Beveridge** (**2005***), but not observed in cryo-electron tomography by ***Beeby et al.** (**2013***)) and cell wall (34 nm according to ***Graham and Beveridge*** (***1994***)). Due to the uncertainty about exact envelope geometry, we thus decided to ignore this effect in our calculations.

### Correction of cytoplasmic mass and surface area for cell-wall thickening during chloramphenical and cerulenin treatment

During chloramphenicol or cerulenin treatments (Figure 3F-G; Figure 4) cell-wall synthesis remains high, and almost unperturbed, according to MreB activity while surface-growth rate drops. Accordingly, cell-wall thickness is expected to increase. However, in our calculation of cytoplasmic surface area ***S*** and mass ***M***, we assume a constant mass fraction of the cell wall (14%) and we implicitly assume a constant distance between the cell contour and the cytoplasmic contour, since our cytoplasmic-contour estimate is based on calibrations with the membrane stain FM4-64 in untreated cells.

To estimate the consequences of these two errors we consider a simple model for corrected cytoplasmic mass and cytoplasmic geometry during excess peptidoglycan synthesis. This then allows us to test and demonstrate that the coordinated increase of surface and mass during chloramphenicol treatment (Figure 3F) remains approximately valid, independently of the approximation, and also if we consider the ratio of cytoplasmic surface and total mass ***M***_tot_ = ***M*** + ***M***_cellwall_ (Figure 3 - figure supplement 6).

For our model correction, we assume that the amount of cell-wall material per surface area and the thickness of the cell wall increase in direct proportion to the difference between normalized cell-wall amount (according to MreB activity) and normalized surface area (both normalized with respect to the unperturbed situation at time=0). This assumption yields a corrected values for ***M***_corr_ = *ζ*(*t*)**M**_tot_, where *ζ*(*t*) now increases in proportion to the amount of peptidoglycan per surface area. Due to the expected thickening of the cell wall, the contour of the cytoplasm is expected to be closer to the cell center by the same absolute amount as the cell wall thickens. Thickening, in turn, is estimated to occur in proportion to *h*(*t*) = [*ζ*(*t*)/ζ_0_]*h*_0_, where *ζ*_0_ = 0.14 (see above), and where *h*_0_ = 34 nm is the height of the unperturbed cell wall (***Graham and Beveridge, 1994***). Cytoplasmic contour correction then leads to a smaller cytoplasmic surface area ***S***_corr_.

For example, peptidoglycan increases by 40% during 20 min after addition of chloramphenicol (since 20 min is half a mass-doubling time in unperturbed cells), while surface increases by about 30% (Figure 3F). Accordingly, the cell wall is expected to increase in mass and thicken by 1.4/1.3 −1 ≈ 8%. The cell-wall mass fraction then increases from 14% to 15.1%, and the cytoplasmic contour is expected to be 2.6 nm smaller (in radial direction) than obtained from phase-contrast microscopy. The comparison between ***S**/**M***, ***S***_corr_/***M***_corr_, and ***S***_corr_/***M***_tot_ (Figure 3 - figure supplement 6) shows that the coupling between the different quantities remains approximately valid independently of the choice of variables. According to similar results based on experiments with vancomycin and cerulenin, cell-wall thickness is expected to change by similar small amounts, which do not affect our interpretation.

### Growth analysis

For bulk growth analysis, cells were cultured in a test tube at 30°C and optical density at 600 nm (OD600) was recorded using a spectrophotometer (Eppendorf). To obtain doubling time, we flt an exponential function to the data points corresponding to the exponential phase (at OD600 between 0.03 and 0.3). For growth analysis from time-lapse microscopy, we calculated relative rates as d(log *X*)/d*t*(*t*_i+0.5_) = 2(*X*_i+1_ – *X*_i_)/(*X*_i+1_ + *X*_i_)/Δ*t*_i_, where ***X*** = ***V***, *S*, *M*, *t*_i+0.5_ = 0.5(*t*_i_ + *t*_i+1_), and Δ*t*_i_ = *t*_i+1_ – *t*_i_. For the display of relative changes of ***V***, ***M***, ***S*** and other quantities, we linearly extrapolated single-cell quantities to *t* = 0 unless stated differently (Figure 5). For display, both relative changes and rates were smoothened with a Gaussian filter with standard deviation of 0.5, if not specified.

### Measurement of MreB motion

We measured MreB motion in two different ways: a) using an epi-fluorescence-based method described below (Method A) and b) a TIRF- and kymograph-based method reported previously (***Dion et al., 2019***) (Method B). The former method is implemented on the same microscope used to conduct all quantitative-phase microscopy. The latter method is implemented on a microscope in Ethan Garner’s lab and was previously demonstrated to give results that agree with high-resolution structured-illumination microscopy (SIM-TIRF) (***Dion et al., 2019***). For epi-fluorescence-based quantification (Method A), we took epi-fluorescence images of MreB-GFP (every 1 s for 30 s) close to the bottom of the cells (about 250 nm below the central plane of cells). Additionally, we took a phase-contrast image to measure the cell contour using the Morphometrics package (***Ursell et al., 2017***) as above. Peak detection and tracking of MreB-GFP were carried out by the Fiji plugin TrackMate(***Tinevez et al., 2017***). Fluorescence spots were detected using the Laplacian of Gaussians (LoG) detector, with a 0.3 μm spot diameter. Tracks were generated using the linear motion LAP tracker, with a search radius 0.15 μm, a minimum displacement of 0.2 μm and 1 frame gap allowed. To quantify the activity of MreB-based cell-wall insertion activity, we measure the sum of all MreB track lengths and divided by total segmented cell area and total observation time (30 s). We refer to this quantity as ‘MreB activity’. If we were able to track all MreB filaments in the field of view, this quantity would be proportional to the areal density of moving filaments times average speed.

For the TIRF-based quantification (Method B), we took images of MreB-GFP by TIRF microscopy (every 300 ms with 1 sec interval for 2 min) followed by a phase-contrast image, and we analyzed the density of directionally moving MreB filaments during cerulenin treatment by the same methods reported by ***Dion et al.** (**2019***). In brief, for every position along the cell centerline, we created a kymograph and subsequently detected moving MreB filaments as described. We counted the directionally moving MreB filaments and then normalized by the projected cell area according to Morphometrics-based segmentation of phase-contrast images obtained at the end of the time-lapse movie to calculate the density of moving MreB fllaments.

### Immersive refractometry

For immersive refractometry we immobilized cells in flow chambers (sticky-slide I Luer 0.1, Ibidi) with a 24×60 mm coverslip (Corning No 1.5) coated with APTES ((3-Aminopropyl)triethoxysilane, Sigma-Aldrich, A3648-100ML): The coverslips were incubated with 2% APTES in ethanol (vol/vol) for 15 min at RT; they were washed with ethanol three times and with distilled water once and then stored in ethanol; before use, ethanol was dried with compressed air. After cell loading we exchanged the media with different refractive index adjusted by Ficoll 400 (Sigma-Aldrich, F4375-100G) and took phase-contrast images. The focal plane was positioned at the middle of the cells.

### Measurement of protein concentration

bAB56 cells were cultured in 50 mL of LB medium at 30°C. Chloramphenicol 100 μg/mL was added to the culture at time = 0 min in Figure 4 - figure supplement 4B. At each time point, cells were harvested from 2 mL of the culture by centrifugation and were suspended in 200 μL of 6 M urea solution. After sonication of the suspension, protein concentration was measured using the Quick Start Bradford Protein Assay (Bio-Rad Laboratories, Inc.)

## Acknowledgements

We thank Richard Wheeler and Ivo Boneca for assistance with cell wall extraction. This work was supported by the European Research Council (ERC) under the Europe Union’s Horizon 2020 research and innovation program [Grant Agreement No. (679980)] to SVT, the French Government’s Investissement d’Avenir program Laboratoire d’Excellence “Integrative Biology of Emerging Infectious Diseases” (ANR-10-LABX-62-IBEID) to SVT, the Mairie de Paris “Emergence(s)” program to SVT, a National Institutes of Health Grant [DP2AI117923-01] to ECG, as well as support from the Volkswagen Foundation to SVT and ECG.

---

**Figure 1 - figure supplement 1.**
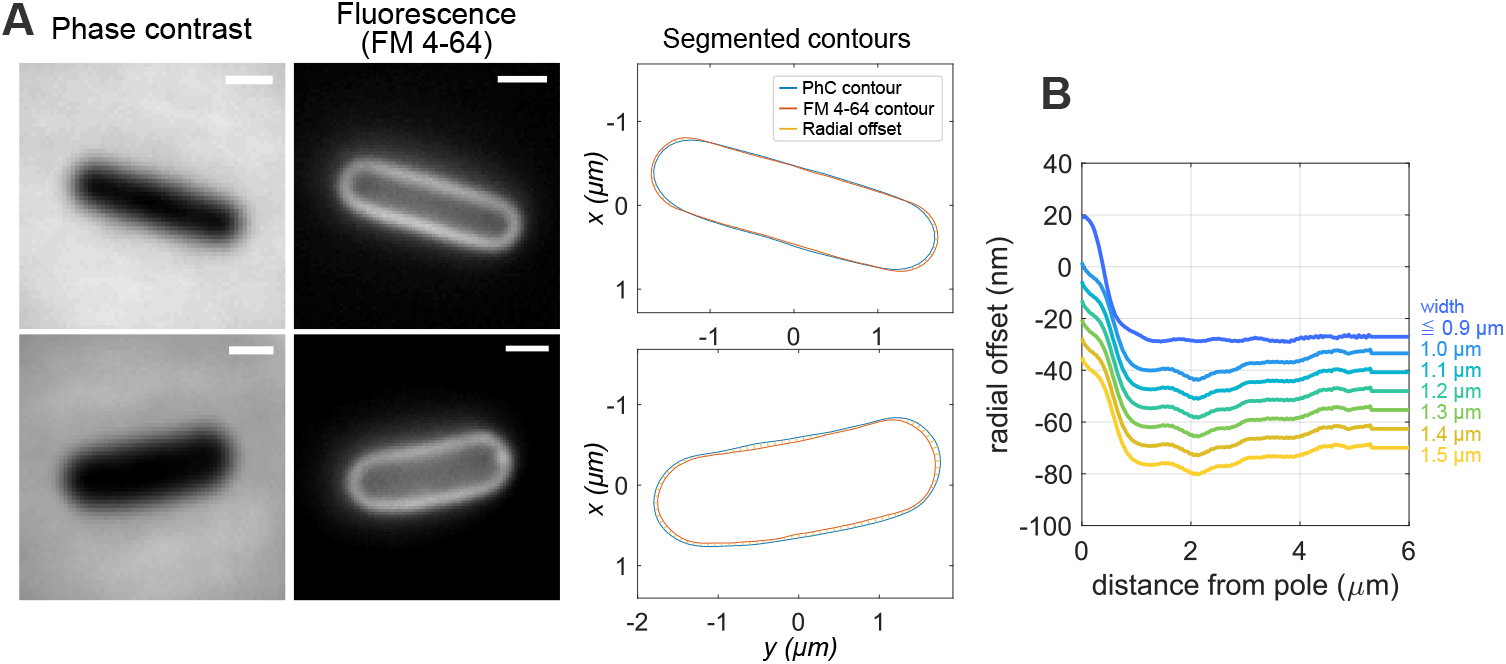
Calibration of phase-contrast contours based membrane contours. **A:** Phase-contrast and fluorescence (membrane stained with FM 4-64) images (**left**) of wild-type cells and bMD586 cells in S7_50_+GlcCaa medium. Membrane contours from the fluorescent images are corrected as described in Materials and Methods. The radial offset is used to correct phase-contrast-based contours for every SLIM image taken during microscopy, conceptually as in ***Oldewurtel et al.** (**2019***). **B:** The comparison for wild-type cells and mutant cells with different *ponA* levels grown in S7_50_+GlcCaa medium (n > 1000 cells) allowed us to infer an average correction as a function of cell width and distance from the cell pole. This function was used to correct the contours of all cells measured with the SLIM module in this study.

**Figure 1 - figure supplement 2.**
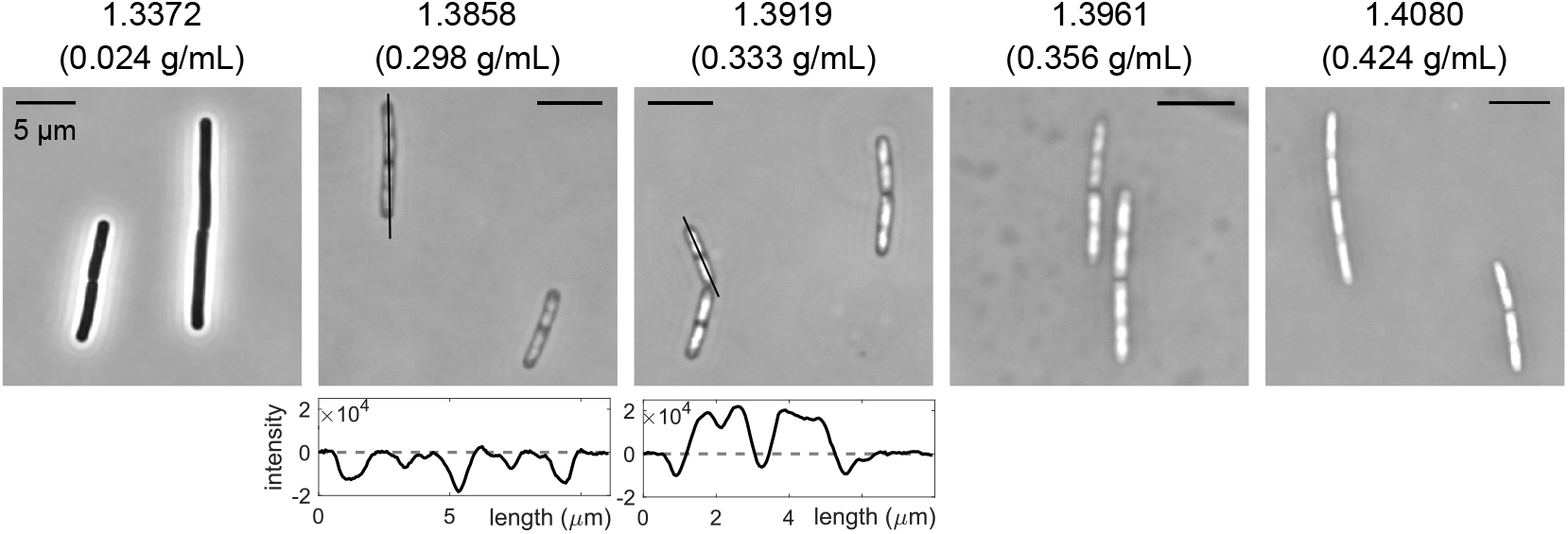
Conf rmation of dry-mass density of cells grown in LB medium by immersive refractometry. **Top:** Phase-contrast-microscopy snaphots of wild-type cells during steady-state growth in LB medium in flow chamber, after attachment to coverslip coated with APTES, with different concentrations of Ficoll 400 that are used to modulate refractive indices (*n* =1.3372-1.4080) as indicated on top of images. **Bottom:** Intensity profiles along lines in phase-contrast images demonstrate spatial heterogeneity of refractive index and thus mass density. If the average refractive index of cells is higher than that of the surrounding medium, the intensity inside cells is lower than outside (observed when *n* =1.3372 or 1.3858). If the refractive index of cells is lower than that of the surrounding medium, the intensity inside cells is higher than outside (observed when *n* =1.3919,1.3961 and 1.4080). Thus, the average refractive cells of cells is in the range of 1.3858-1.3919, which corresponds to a mass-density range of 0.298-0.333 g/mL, compatible with Figure 1B.

**Figure 1 - figure supplement 3.**
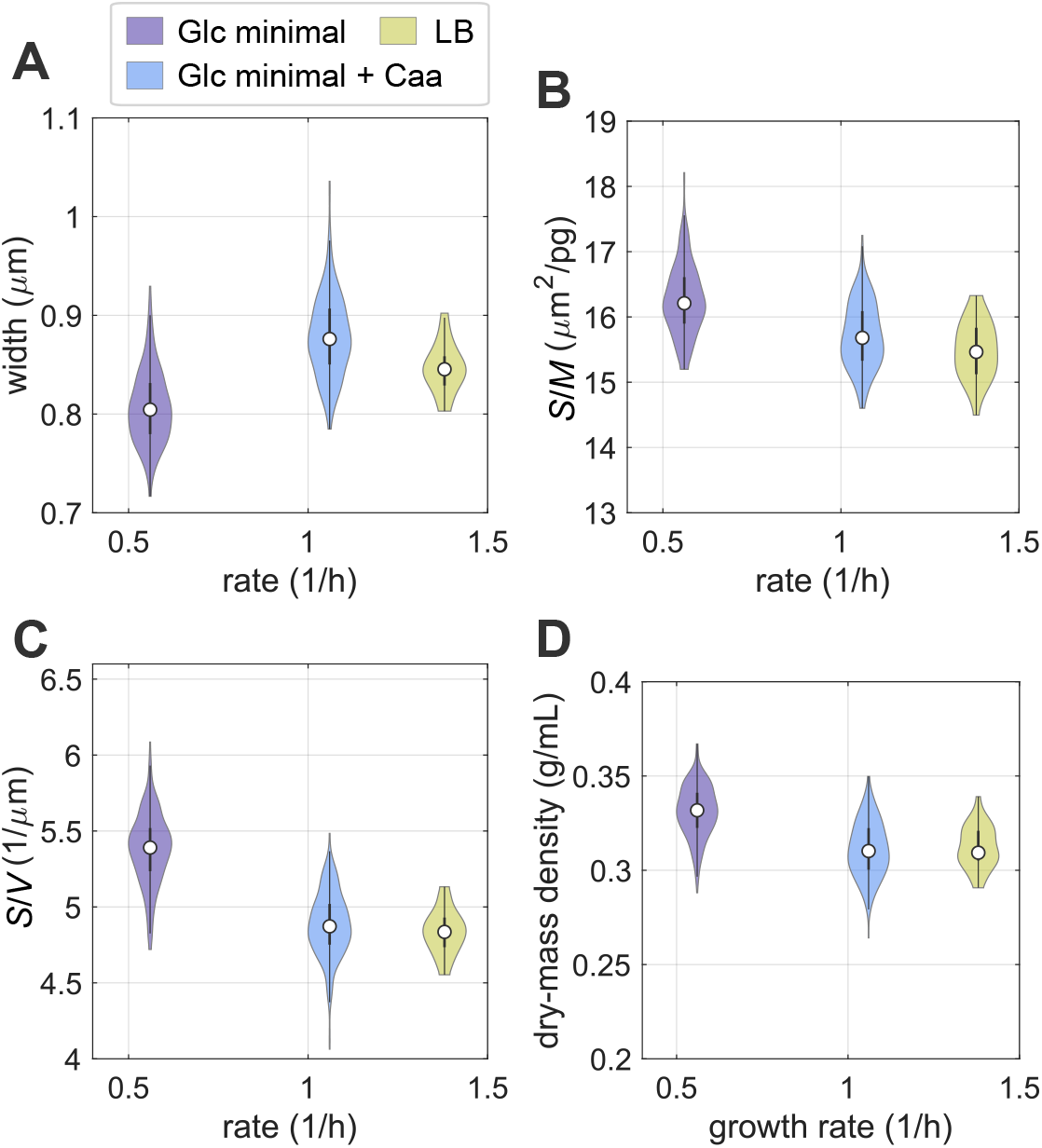
Single-cell properties during steady-state growth in different nutrient conditions. Width (**A**), surface-to-mass ratio (**B**), surface-to-volume ratio (**C**), and dry-mass density (**D**) of wild-type cells cultured in S7_50_+Glc, and S7_50_+GlcCaa, and LB medium at 30°C. The same dataset presented in Figure 1B. (white circles = median; grey rectangles = interquartile range).

**Figure 1 - figure supplement 4.**
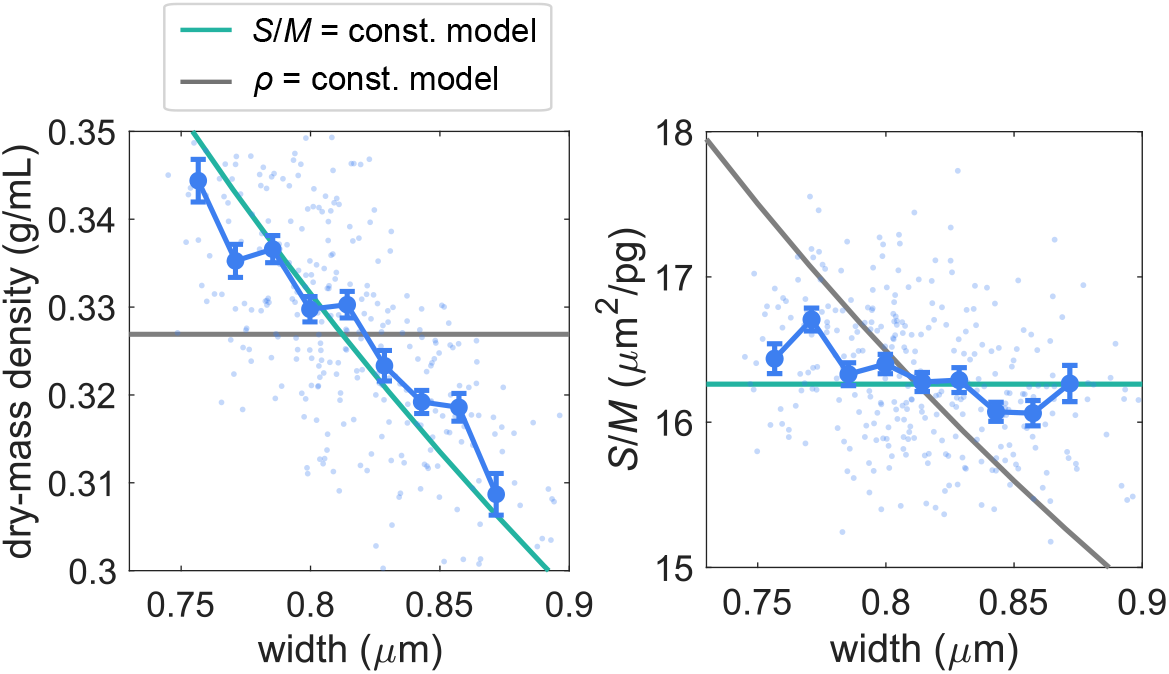
Width dependency of dry-mass density and surface-to-mass ratio of wild-type cells during steady-state growth in S7_50_+Glc. Blue dots: values of single wild-type cells; blue symbols and line: binned averages ± SE; green lines: model prediction for spherocylinder with constant surface-to-mass ratio; gray lines: model prediction for spherocylinder with constant dry-mass density.

**Figure 1 - figure supplement 5.**
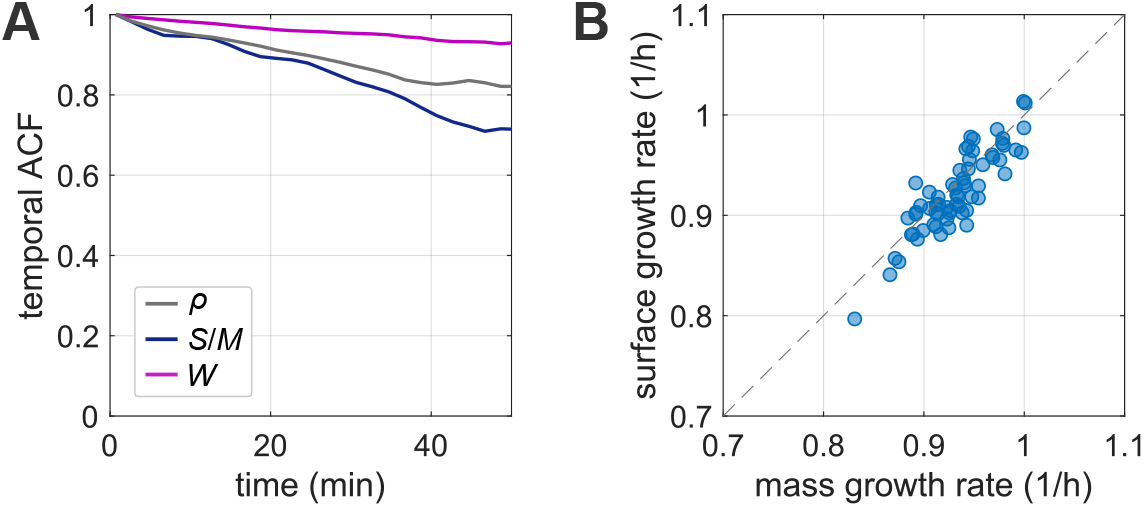
Autocorrelation functions of width, surface-to-mass ratio and mass density, as well as surface- versus mass-growth rates during steady-state growth. **A:** Temporal autocorrelation function (ACF) of width, surface-to-mass ratio and mass density during steady-state growth. Here, we define the ACF as 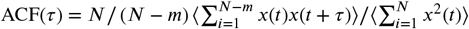, where *τ* = *m*Δ*t* with Δ*t* the time step. *x* = *X* – 〈*X*〉 are the normalized single-cell quantities (*X* = *ρ*, ***S**/**M***, ***W***). Angular brackets denote an average over all cells. **B:** Single-cell mass growth rate ***λ_M_*** and surface growth rate *λ_S_* are tightly correlated during steady-state growth. Here, to reduce measurement noise, we smoothened mass and surface by a Gaussian fllter with standard deviation of 7Δ*t* (Δ*t* = 2 s) and subsequently smoothened growth rates with the same fllter. While variations of *λ_S_* and *λ_**M**_* are each about 0.04 (CV), the standard deviation of λ*_**S**_* - λ_***M***_ is about 0.02〈λ_***M***_〉. Dots: single-cell measurements, dashed line: identity function.

**Figure 1 - figure supplement 6.**
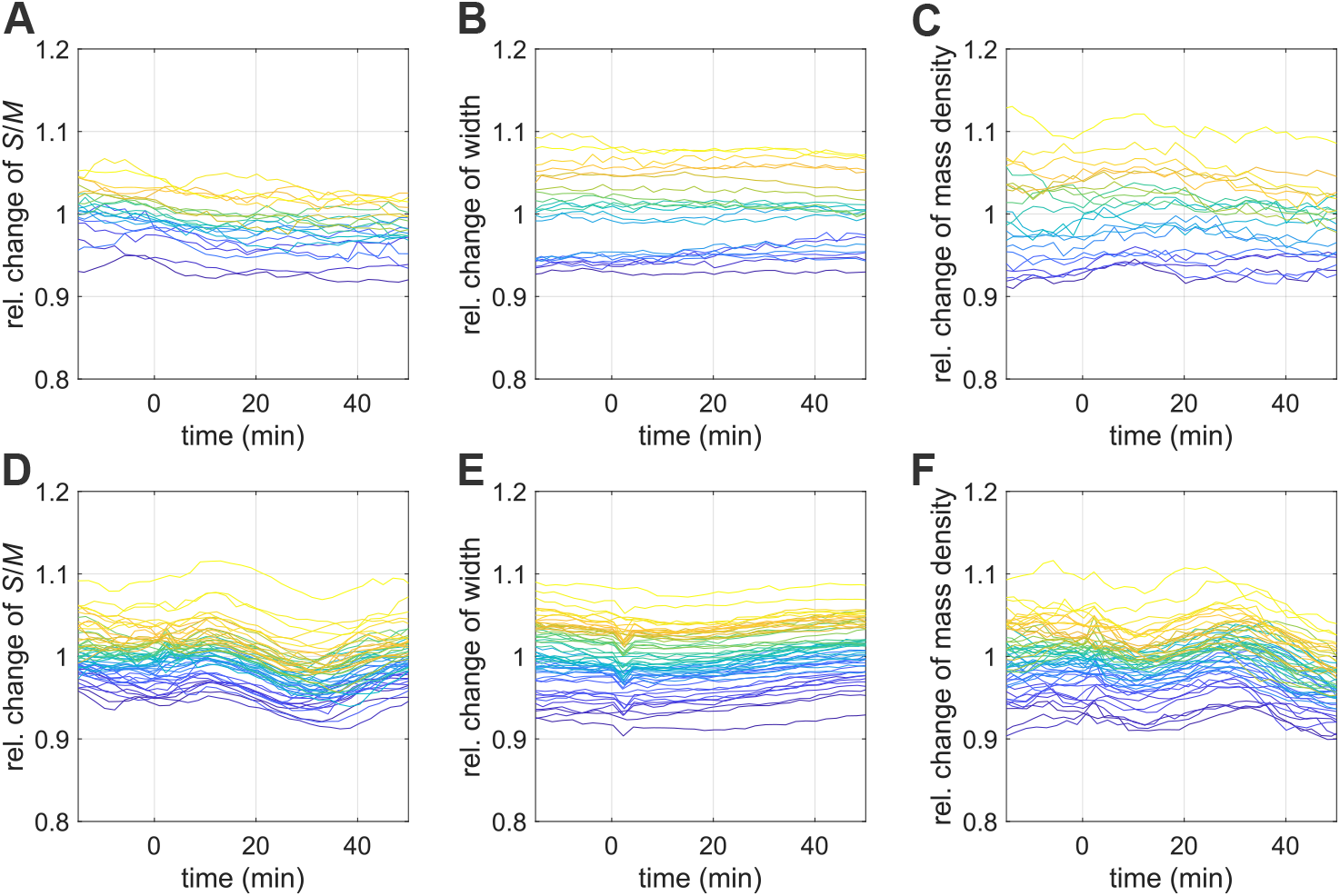
Single-cell traces during nutrient shifts. Relative changes of surface-to-mass ratio (**A**), width (**B**), dry-mass density (**C**) during nutrient upshift (the same experiment shown in Figure 1E). Relative surface-to-mass ratio (**D**), width (**E**), dry-mass density (**F**) during nutrient downshift (the same experiment shown in Figure 1F).

**Figure 2 - figure supplement 1.**
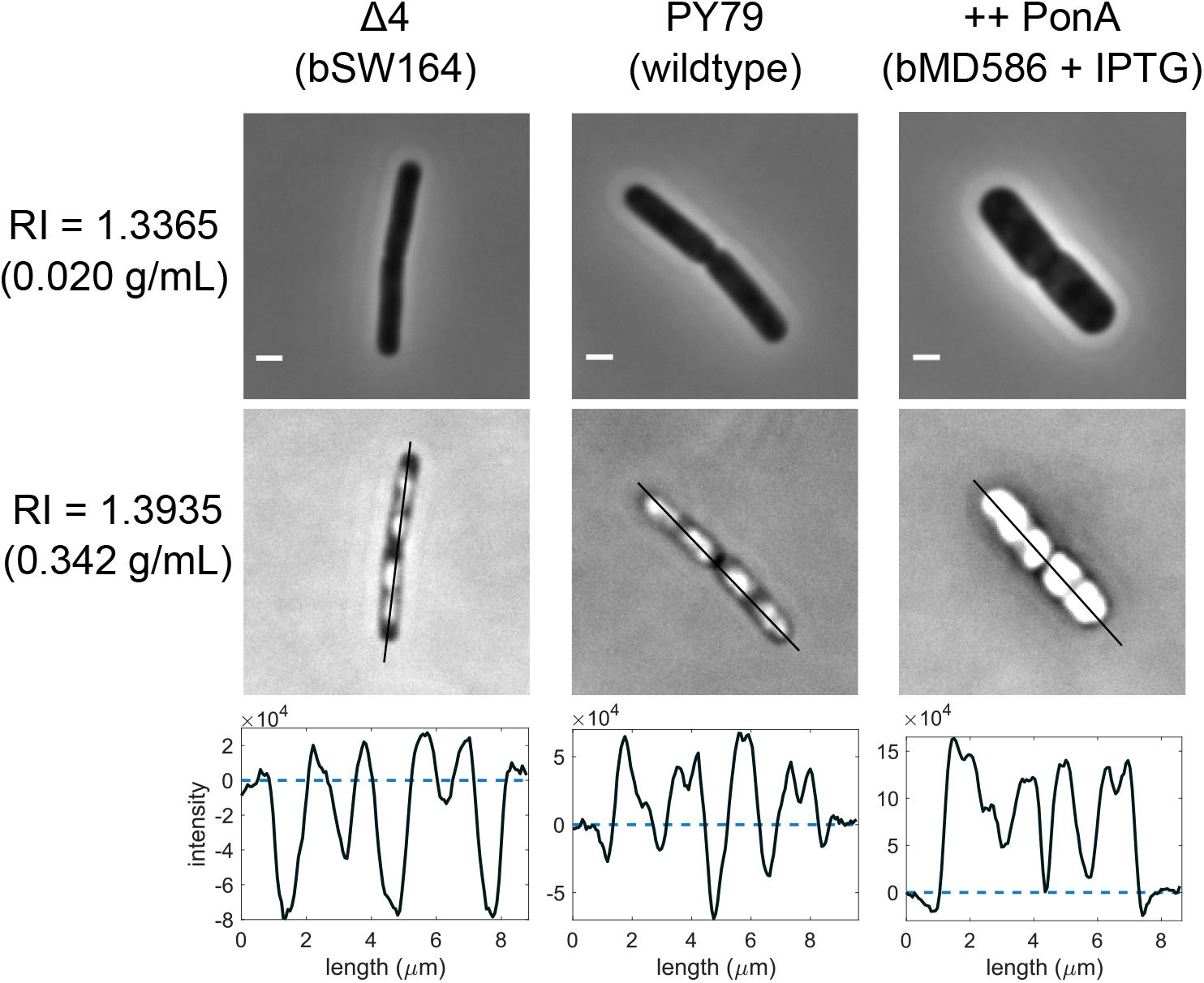
Immersive refractometry of wildtype and mutants with different aPBP-expression levels. Snapshots of wild-type, bSW164 and bMD586 cells after steady-state growth in S7_50_+GlcCaa medium immobilized in flow chamber. To overexpress PonA, bMD586 cells was cultured with 1 mM IPTG. **Top:** Snapshots of cells in flow chamber filled with media of different refractive indices due to supplementation with Ficoll 400 (*n* =1.3365,1.3935) taken by phase-contrast microscopy (for an explanation see also Figure 1 - figure supplement 2). **Bottom:** Intensity profiles along lines in top panels.

**Figure 2 - figure supplement 2.**
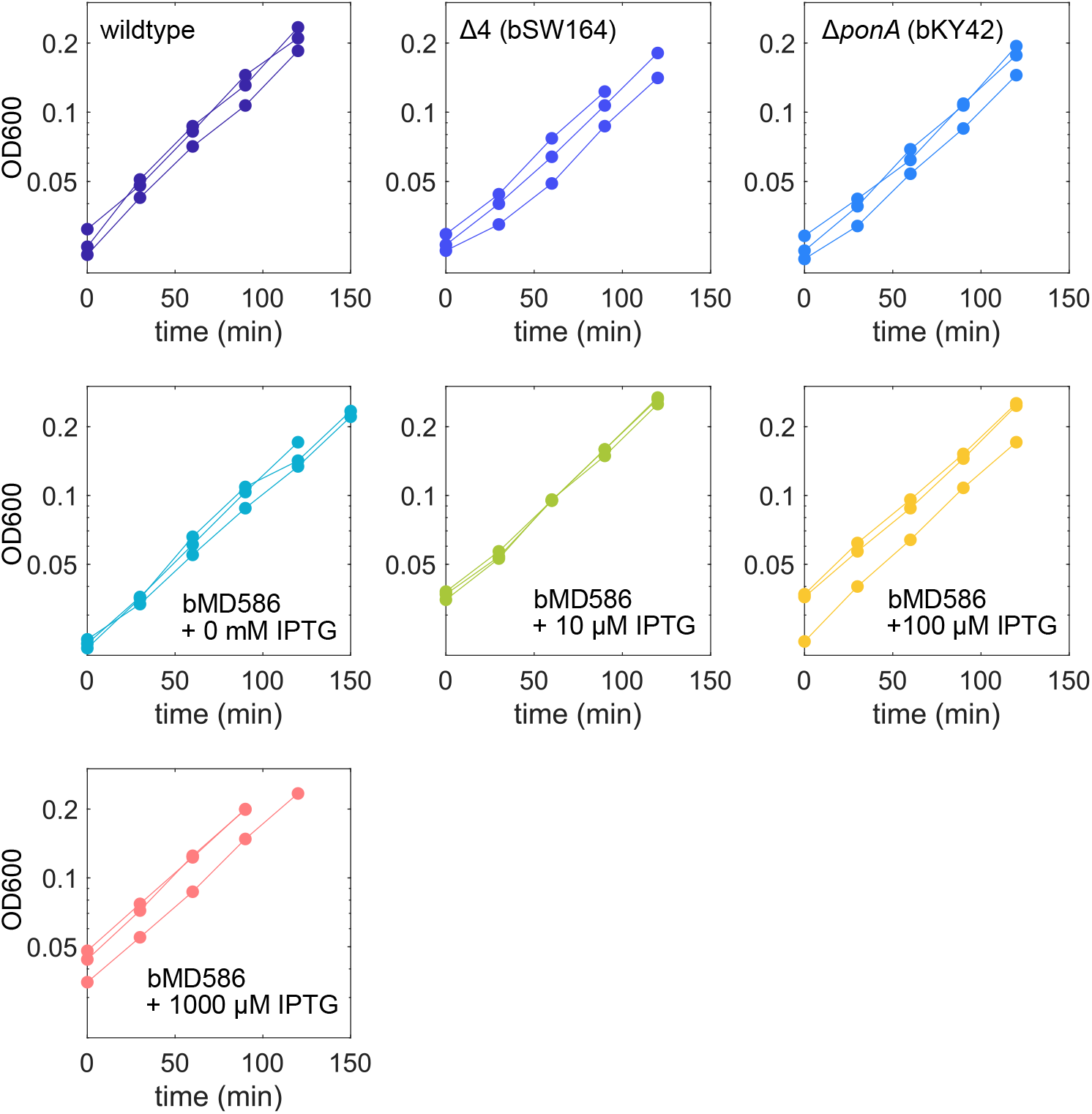
Growth curves of cells with different expression level of aPBPs. Optical density (OD600) as a function of time for the same experiment shown in Figure 2F. Wild-type, bSW164, bKY42, and bMD586 cells were cultured in S7_50_+GlcCaa medium. To overexpress PonA, 10-1000 μM of IPTG was added to bMD586 cultures. Every line represents an independent biological replicate.

**Figure 3 - figure supplement 1.**
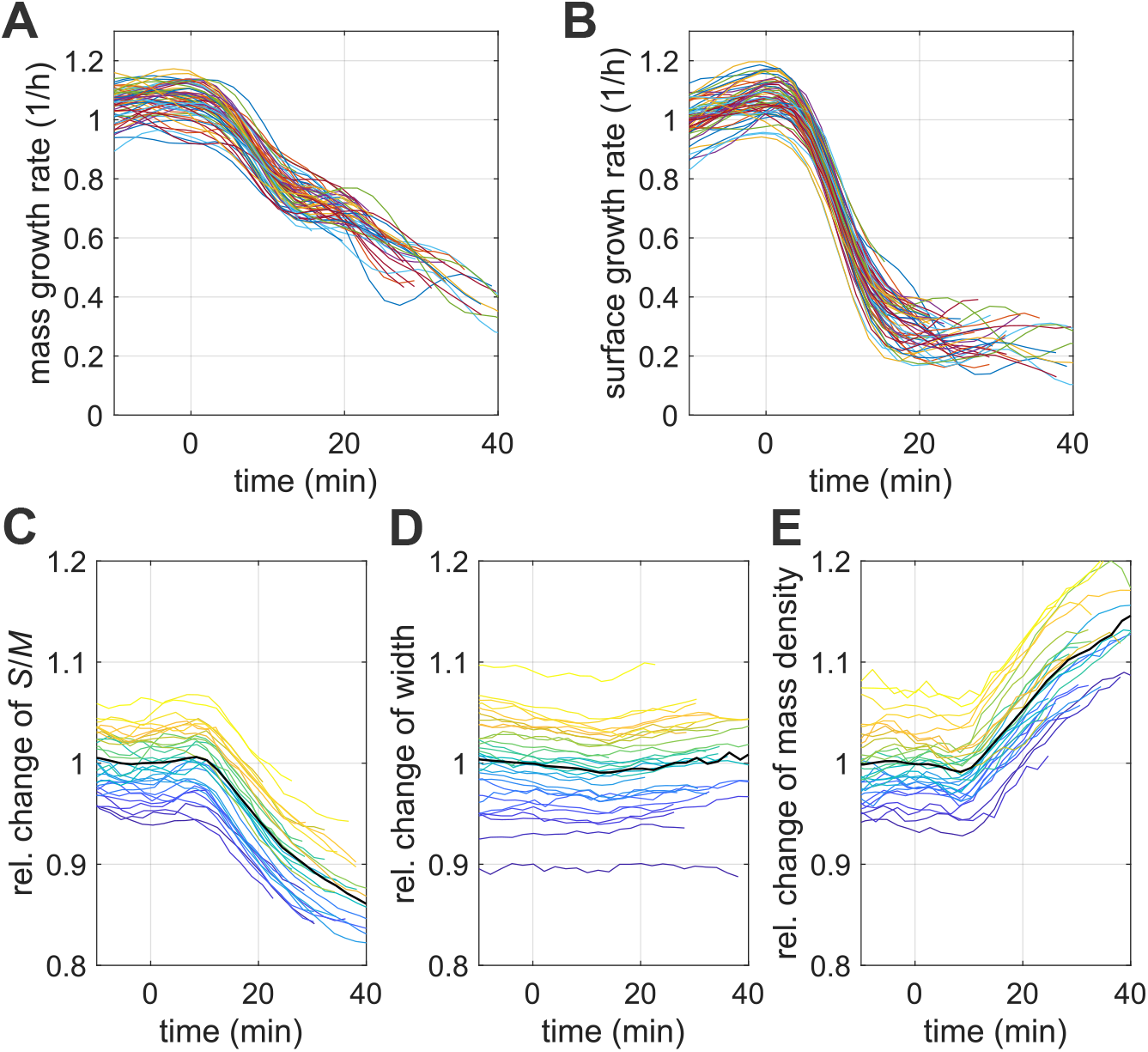
Single-cell behavior during vancomycin treatment. Growth rate of mass (**A**) and surface (**B**), relative changes of surface-to-mass ratio (**C**), width (**D**), dry-mass density (**E**) during vancomycin treatment. (the same experiment of Figure 3A, B). For better visibility, growth rates were smoothened with a Gaussian fllter with standard deviation of 2Δ*t* where Δ*t* = 2 min is the time interval.

**Figure 3 - figure supplement 2.**
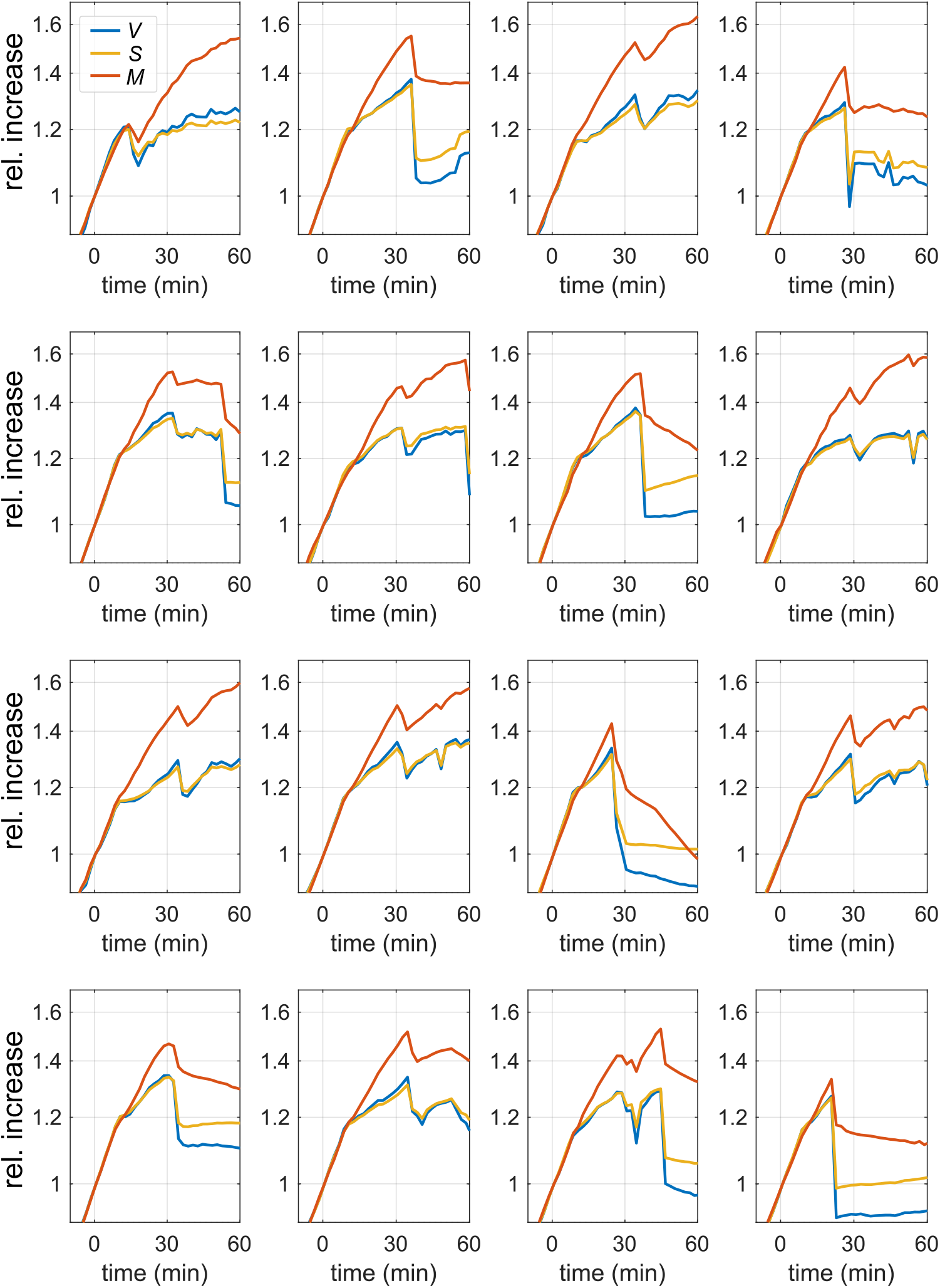
Single cells lose parts of their mass during vancomycin treatment. Relative increase of volume, surface and dry mass of single cells that showed a transient reduction of more the 2% of dry mass during vancomycin treatment (the same experiment of Figure 3A, B).

**Figure 3 - figure supplement 3.**
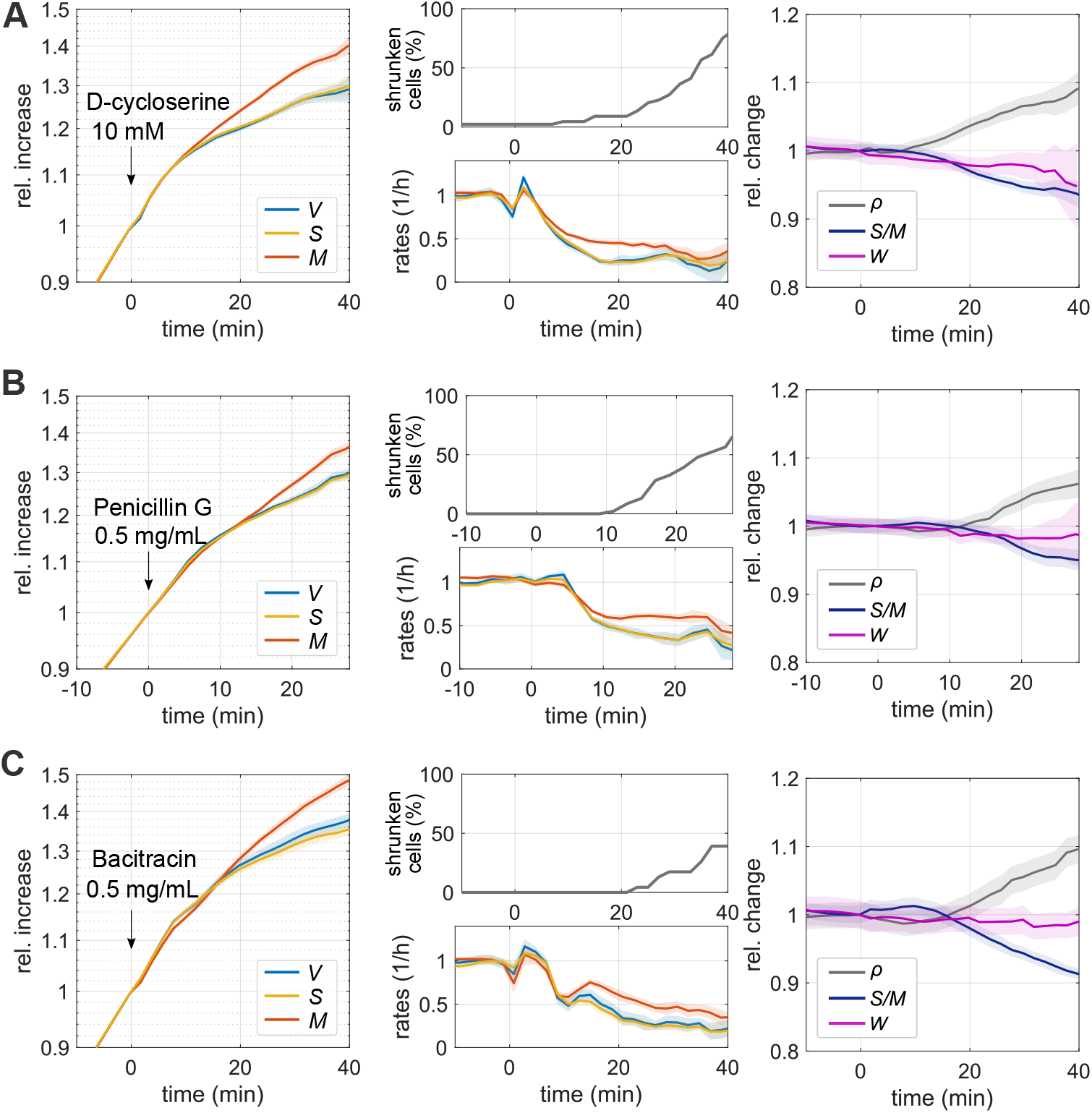
Time lapses of single cells during inhibition of peptidoglycan synthesis using different drugs. **A:** Single-cell time lapse of bAB56 cells (fllamenting mutant) treated with D-cycloserine (10 mM). The drug was added on top of the agarose pad (S7_50_+GlcCaa) about 30 min after placing cells on the microscope (at time=0). To avoid cell division, MciZ was induced 30 min prior to microscopy. Relative increase (left) and rates (middle-bottom) of volume, surface and dry mass. Lysis rates (middle-top). Relative change of dry-mass density, surface-to-mass ratio, and width (right). (Solid lines + shadings = average ± SE) **B:** Single-cell time lapse of bAB56 cells (fllamenting mutant) treated with Penicillin G (0.5 mg/mL). Otherwise the same as A. **C:** Single-cell time lapse of bAB56 cells (fllamenting mutant) treated with Bacitracin (0.5 mg/mL). Otherwise the same as A.

**Figure 3 - figure supplement 4.**
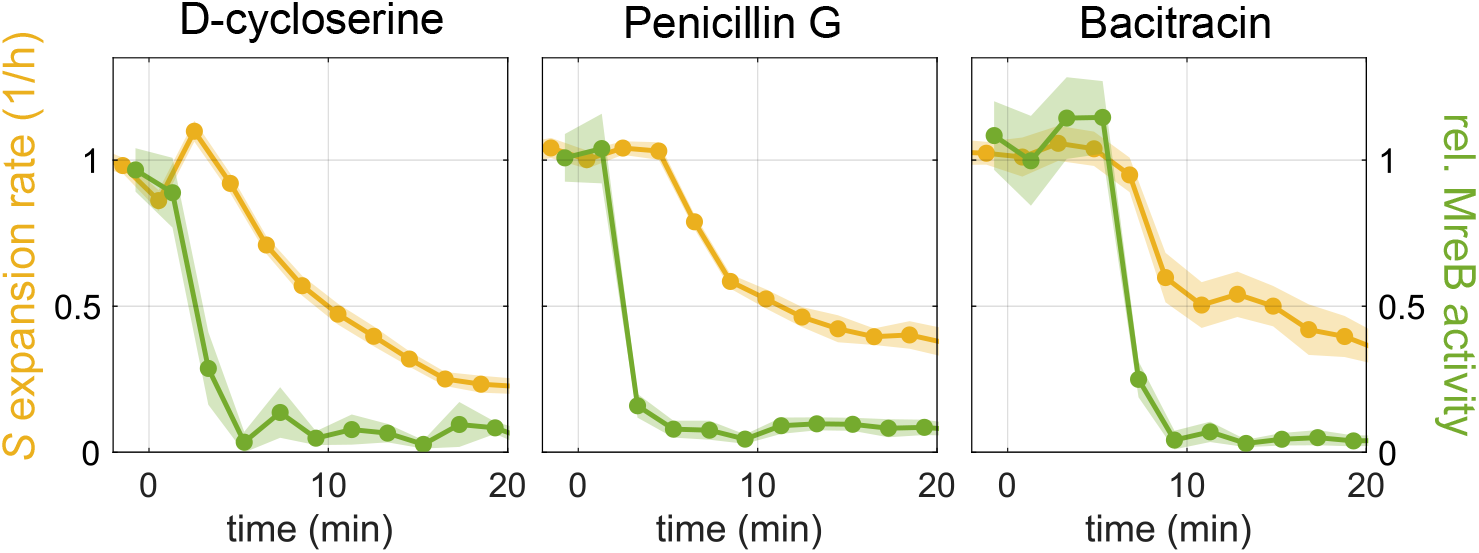
Surface-expansion rate and MreB activity during inhibition of peptidoglycan synthesis using different drugs. Relative surface-expansion rate obtained from Figure 3 - figure supplement 3 (yellow, left axis) and relative MreB activity (green, right axis), inferred from MreB-GFP movies of bYS19 cells as described for Figures 3, 4, during treatment with D-cycloserine 10 mM, Penicillin G 0.5 mg/mL, or Bacitracin 0.5 mg/mL in the same way as Figure3 - figure supplement 3 (Solid lines + shadings = average ± SE). MreB activity was quantified as described in Materials and Methods.

**Figure 3 - figure supplement 5.**
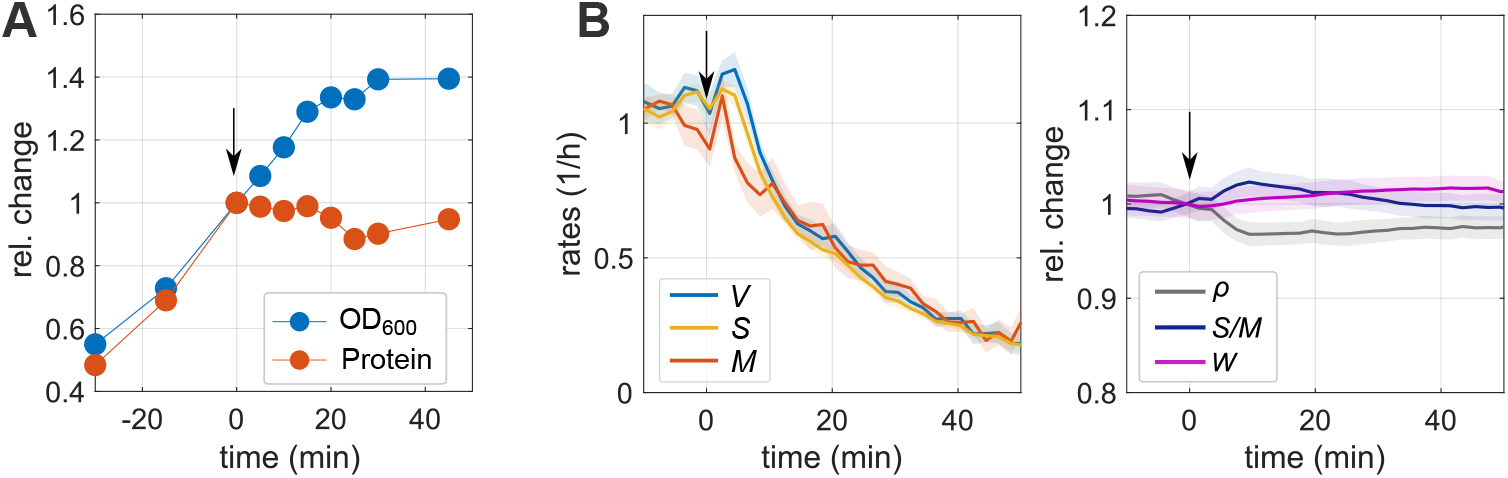
Chloramphenicol treatment arrests protein synthesis but does not arrest surface growth. **A:** Relative increase of optical density (OD600) and protein amount according to Quick Start Bradford assay during chloramphenicol treatment. Wild-type cells were cultured in LB medium, and 100 μg/mL of chloramphenicol was added at time=0. **B:** Single-cell time lapse of filamenting cells (bAB56) cultured in S7_50_+GlcCaa medium and treated with chloramphenicol 100 μg/mL at time=0. (The same experiment shown in Figure 3F). Chloramphenicol was added on top of the agarose pad (S7_50_+GlcCaa) about 30 min after placing cells on the microscope. To avoid cell division, MciZwas induced 30 min prior to microscopy. **Left:** Rates of volume, surface and dry mass. **Right:** Relative change of dry-mass density, surface-to-mass ratio, and width. (Solid lines + shadings = average ± SE)

**Figure 3 - figure supplement 6.**
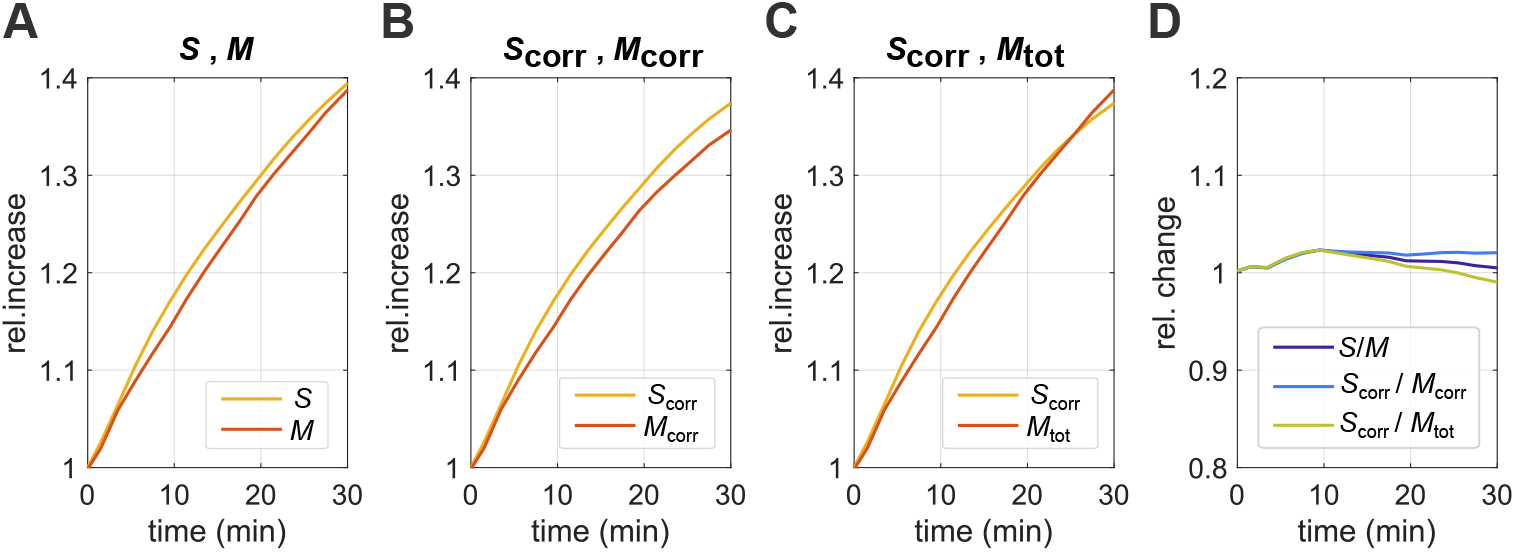
Considering potential effects of continued elevated cell-wall synthesis during chloramphenicol treatment on the surface-to-mass coupling. Relative increase of surface and mass of chloramphenicol-treated cells as in Figure 3F-G, before cell-wall thickening is taken into account (**A**), if both ***S*** and ***M*** are corrected for possible effects of cell-wall thickening (**B**; see Materials and Methods for the correction) or if we consider the corrected surface area and the total cell mass *M*_tot_ = ***M*** + ***M***_cellwall_ (**C**). **D:** Relative changes of the different surface-to-mass ratios, ***S***_corr_/***M***_corr_, ***S***_corr_/***M***_tot_ show small variations if compared to our main method. Thus, a correction of ***S*** and ***M*** does not alter our conclusion that ***S*** and ***M*** remain coupled during chloramphenicol treatment.

**Figure 4 - figure supplement 1.**
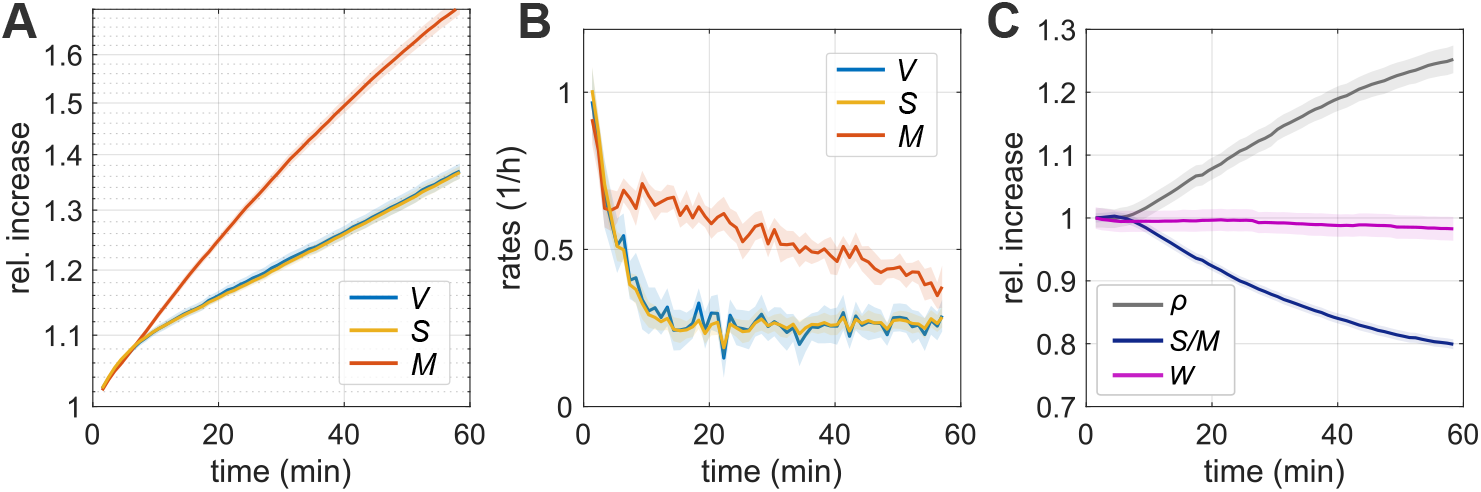
Single-cell time lapse during treatment with cerulenin contained in the agarose pad prior to microscopy. **A-C:** Single-cell time lapse of bAB56 cells (fllamenting mutant) treated with cerulenin 100 μg/mL. Cerulenin was contained in the agarose pad (S7_50_+GlcCaa) so that cells are immediately exposed to the drug at its final concentration. To avoid cell division, MciZ was induced 30 min prior to microscopy. We took images every 1 min. Relative increase (**A**) and rates (**B**) of volume, surface and dry mass. (**C**) Relative change of dry-mass density, surface-to-mass ratio, and width. (Solid lines + shadings = average ± SE)

**Figure 4 - figure supplement 2.**
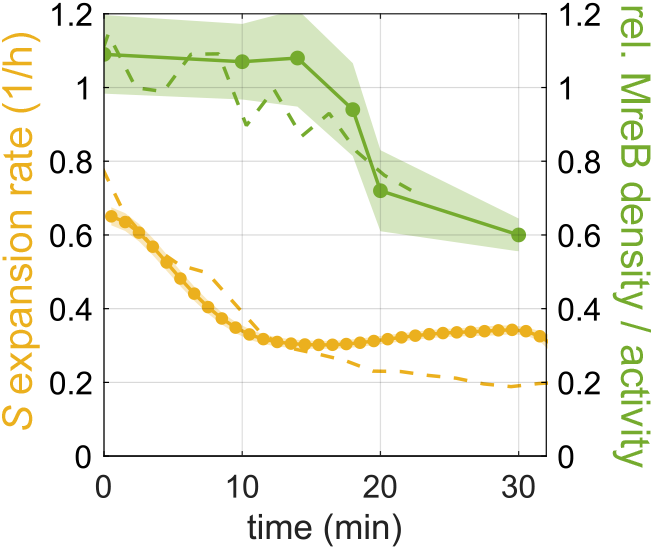
Complementary method to measure MreB-based cell-wall insertion after cerulenin treatment. Solid lines: Surface expansion rate (yellow) and relative density of directionally moving MreB fllaments (green) of bYS19 cells treated with cerulenin 100 μg/mL (Solid lines + shadings = average ± SE). Different from Figure 4, cerulenin was contained in the agarose pad (S7_50_+GlcCaa). Density of directionally moving MreB filaments was measured based on TIRF-imaging and the analysis of kymographs as previously reported (***Dion et al., 2019***). Surface expansion rate was calculated based on time-lapse movies with 1 min interval. Dashed lines: For comparison, we also indicated surface expansion rate and relative MreB activity based on epi-fluorescence movies already presented in Figure 4, with the time shifted by 7 min as an estimated time when cells are exposed to the minimal inhibitory concentration (***Schujman et al., 2001***) according to a 1-dimensional diffusion equation.

**Figure 4 - figure supplement 3.**
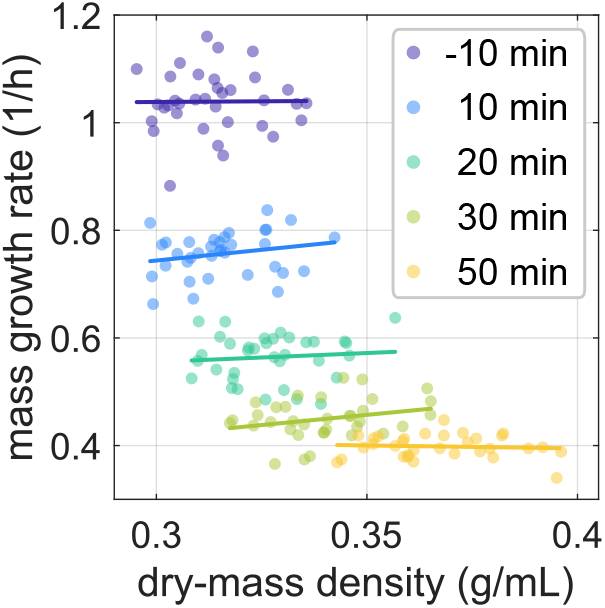
Relationship between dry-mass density and mass growth rate during cerulenin treatment. Single-cell mass growth rate shows no visible correlation with single-cell dry-mass density at different times after cerulenin treatment (same experiment shown in Figure 4A). Dots: single-cell measurements, lines: linear regression. Mass growth rates were smoothened with a Gaussian filter with standard deviation of 2Δ*t*, where Δ*t* = 2 min is the time interval.

**Figure 4 - figure supplement 4.**
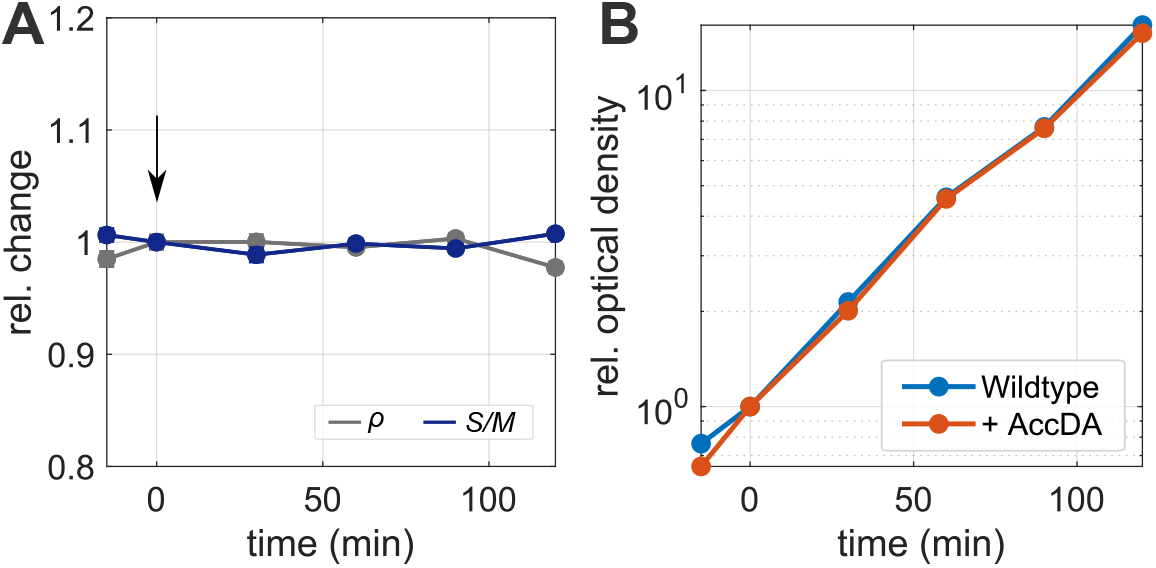
Control experiment for AccDA overexpression. **A:** Relative change of dry-mass density and surface-to-mass ratio obtained from snapshots of wild-type cells (average ± SE) cultured in liquid LB medium. Xylose 10 mM was added at time = 0 min. **B:** Relative optical density (OD600) during the experiment in (A) and shown in Figure 4D. Values are corrected for backdilution, to maintained OD below 0.3.

**Figure 5 - figure supplement 1.**
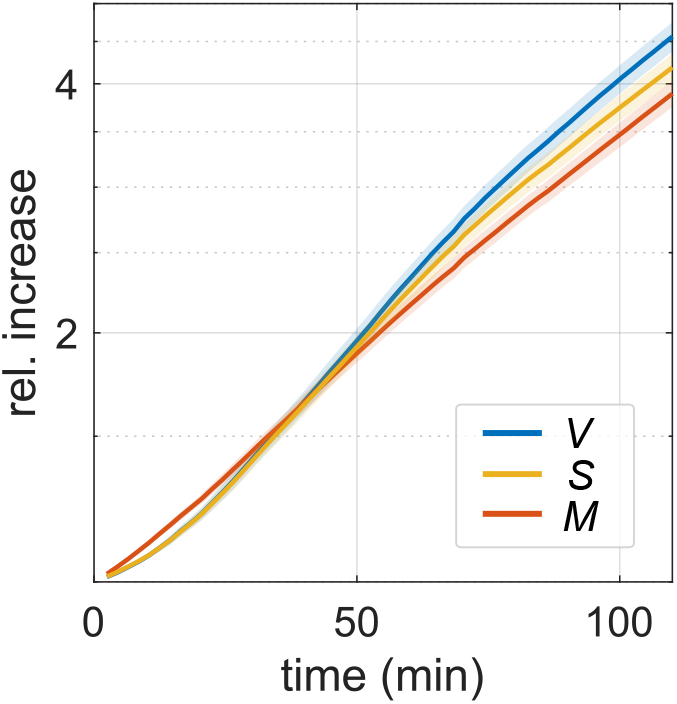
Growth of bAB56 cells during recovery from cerulenin treatment. The same experiment shown in Figure 5 A-B. Relative increase of volume, surface and dry mass.(Solid lines + shadings = average ± SE)

**Figure 5 - figure supplement 2.**
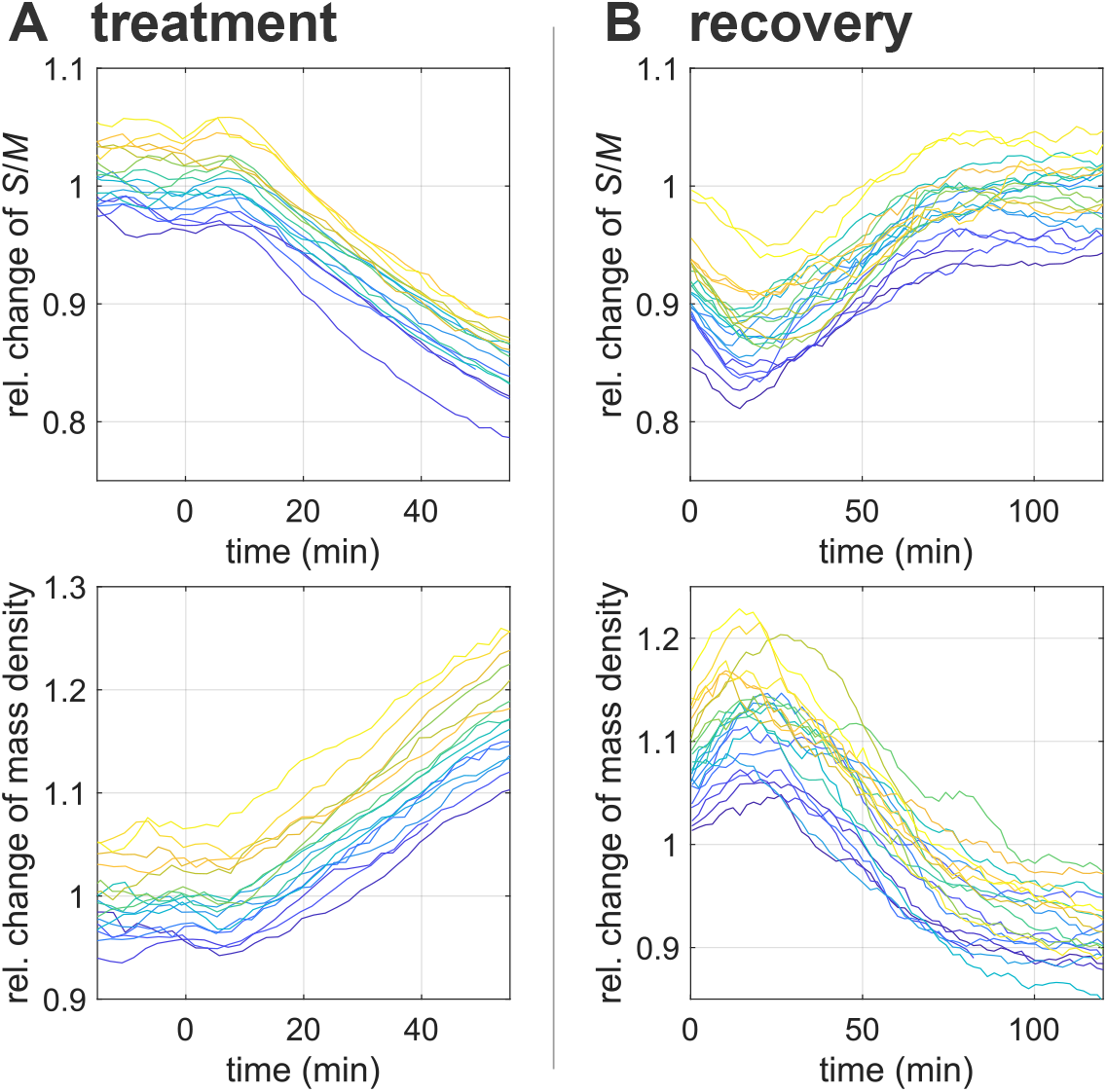
Single-cell traces during cerulenin treatment and recovery. Single-cell changes of surface-to-mass ratio (**top**) and dry-mass density (**bottom**) during cerulenin treatment (**A**) (the same experiment shown in Figure 4A) and during recovery from 30 min-long cerulenin treatment in bulk (**B**) (the same experiment shown in Figure 5A-B) after normalization with respect to the respective average steady-state (unperturbed) values.

## Supplementary files

**Supplementary File 1.** Detailed information of snapshot-,time-lapse-,and MreB-experiments. We provide the following information: corresponding figures; strain used; growth medium; conversion factors used to normalize width, surface-to-mass ratio, mass density, MreB activity in plots depicting relative changes; number of considered cells; time of MciZ induction prior to placing cells on agarose pad.

**Supplementary File 2.** Oligonucleotides, DNA fragments and strains used in this study.

**Supplementary File 3.** Chemical compositions of *B. subtilis* cell and their refraction increments.

## Supplementary Videos

**Figure 1 - video 1.** Phase-contrast microscopy of single fllamenting cell (bAB56) during nutrient upshift.

Corresponds to Fig 1E.

**Figure 1 - video 2.** Phase-contrast microscopy of single fllamenting cell (bAB56) during nutrient downshift. Corresponds to Fig 1F.

**Figure 3 - video 1.** Phase-contrast microscopy of single fllamenting cell (bAB56) during vancomycin treatment. Corresponds to Fig 3A-B,D.

**Figure 3 - video 2.** MreB-GFP rotation in single cell (bYS19) before vancomycin treatment. Images were acquired for 30 s at a rate of 1/s. Corresponds to Fig 3C-D.

**Figure 3 - video 3.** MreB-GFP rotation in single cell (bYS19) after vancomycin treatment. Corresponds to Fig 3C-D.

**Figure 3 - video 4.** Phase-contrast microscopy of single fllamenting cell (bAB56) during chloramphenicol treatment. Corresponds to Fig 3F-G.

**Figure 3 - video 5.** MreB-GFP rotation in single cell (bYS19) after chloramphenicol treatment. Corresponds to Fig 3G.

**Figure 4 - video 1.** Phase-contrast microscopy of single fllamenting cell (bAB56) during cerulenin treatment. Corresponds to Fig 4A, C.

**Figure 4 - video 2.** MreB-GFP rotation in single cell (bYS19) after cerulenin treatment. Corresponds to Fig 4B, C.

**Figure 5 - video 1.** Phase-contrast microscopy of single fllamenting cell (bAB56) during recovery from cerulenin treatment. Corresponds to Fig 5A, B.

## Notes

### Competing Interest Statement

The authors have declared no competing interest.

## References

Barer R. Phase contrast and interference microscopy in cytology. Physical techniques in biological research. 1956; 3:29–90.

Barer R, Joseph S. Refractometry of living cells: Part I. Basic principles. Journal of Cell Science. 1954; 3(32):399–423.

Barna J, Williams D. The structure and mode of action of glycopeptide antibiotics of the vancomycin group. Annual review of microbiology. 1984; 38(1):339–357.

Beeby M, Gumbart JC, Roux B, Jensen GJ. Architecture and assembly of the G ram-positive cell wall. Molecular microbiology. 2013; 88(4):664–672.

Billaudeau C, Chastanet A, Yao Z, Cornilleau C, Mirouze N, Fromion V, Carballido-López R. Contrasting mechanisms of growth in two model rod-shaped bacteria. Nature communications. 2017; 8(1):1–11.

Bishop D, Rutberg L, Samuelsson B. The chemical composition of the cytoplasmic membrane of Bacillus subtilis. European journal of biochemistry. 1967; 2(4):448–453.

Carballido-López R, Formstone A, Li Y, Ehrlich SD, Noirot P, Errington J. Actin homolog MreBH governs cell morphogenesis by localization of the cell wall hydrolase LytE. Developmental cell. 2006; 11(3):399–409.

Cayley S, Lewis BA, Guttman HJ, Record Jr MT. Characterization of the cytoplasm of Escherichia coli K-12 as a function of external osmolarity: implications for protein-DNA interactions in vivo. Journal of molecular biology. 1991; 222(2):281–300.

Chung K. Autoradiographic studies of bacterial cell wall replication: I. Cell wall growth of Bacillus cereus in the presence of chloramphenicol. Canadian journal of microbiology. 1967; 13(4):341–350.

Cronan Jr JE, Rock CO. Biosynthesis of membrane lipids. EcoSal Plus. 2008; 3(1).

Cronan Jr JE, Waldrop GL. Multi-subunit acetyl-CoA carboxylases. Progress in lipid research. 2002; 41(5):407–435.

D’agnony G, Rosenfeld IS, Awaya J, Omura S, Vagelos PR. Inhibition of fatty acid synthesis by the antibiotic cerulenin: Specific inactivation of *β*-ketoacyl-acyl carrier protein synthetase. Biochimica et Biophysica Acta (BBA)-Lipids and Lipid Metabolism. 1973; 326(2):155–166.

Daniel RA, Errington J. Control of cell morphogenesis in bacteria: two distinct ways to make a rod-shaped cell. Cell. 2003; 113(6):767–776.

Delarue M, Brittingham G, Pfeffer S, Surovtsev I, Pinglay S, Kennedy K, Schaffer M, Gutierrez J, Sang D, Poterewicz G, et al. mTORC1 controls phase separation and the biophysical properties of the cytoplasm by tuning crowding. Cell. 2018; 174(2):338–349.

Dion MF, Kapoor M, Sun Y Wilson S, Ryan J, Vigouroux A, Van Teeffelen S, Oldenbourg R, Garner EC. Bacillus subtilis cell diameter is determined by the opposing actions of two distinct cell wall synthetic systems. Nature microbiology. 2019; 4(8):1294–1305.

Domínguez-Cuevas P, Porcelli I, Daniel RA, Errington J. Differentiated roles for MreB-actin isologues and autolytic enzymes in B acillus subtilis morphogenesis. Molecular microbiology. 2013; 89(6):1084–1098.

Domínguez-Escobar J, Chastanet A, Crevenna AH, Fromion V, Wedlich-Söldner R, Carballido-López R. Processive movement of MreB-associated cell wall biosynthetic complexes in bacteria. Science. 2011; 333(6039):225–228.

Dourado H, Lercher MJ. An analytical theory of balanced cellular growth. Nature communications. 2020; 11(1):1–14.

Dowhan W. A retrospective: use of Escherichia coli as a vehicle to study phospholipid synthesis and function. Biochimica et Biophysica Acta (BBA)-Molecular and Cell Biology of Lipids. 2013; 1831(3):471–494.

Edelstein A, Amodaj N, Hoover K, Vale R, Stuurman N. 933 Computer control of microscopes using microManager. Current protocols. 2010; 934.

Freese E, Klofat W, Galliers E. Commitment to sporulation and induction of glucose-phosphoenolpyruvate-transferase. Biochimica et Biophysica Acta (BBA)-General Subjects. 1970; 222(2):265–289.

Garner EC, Bernard R, Wang W, Zhuang X, Rudner DZ, Mitchison T. Coupled, circumferential motions of the cell wall synthesis machinery and MreB fllaments in B. subtilis. Science. 2011; 333(6039):222–225.

Graham L, Beveridge T. Structural differentiation of the Bacillus subtilis 168 cell wall. Journal of bacteriology. 1994; 176(5):1413–1421.

Handler AA, Lim JE, Losick R. Peptide inhibitor of cytokinesis during sporulation in Bacillus subtilis. Molecular microbiology. 2008; 68(3):588–599.

Höltje JV. Growth of the stress-bearing and shape-maintaining murein sacculus of Escherichia coli. Microbiology and molecular biology reviews. 1998; 62(1):181–203.

Huff E. Lipoteichoic acid, a major amphiphile of Gram-positive bacteria that is not readily extractable. Journal of bacteriology. 1982; 149(1):399.

Hussain S, Wivagg CN, Szwedziak P Wong F Schaefer K, Izoré T Renner LD, Holmes MJ, Sun Y Bisson-Filho AW, et al. MreB fllaments align along greatest principal membrane curvature to orient cell wall synthesis. Elife. 2018; 7:e32471.

Jaacks K, Healy J, Losick R, Grossman A. Identification and characterization of genes controlled bythe sporulation-regulatory gene spo0H in Bacillus subtilis. Journal of bacteriology. 1989; 171(8):4121–4129.

Kawai Y, Asai K, Errington J. Partial functional redundancy of MreB isoforms, MreB, Mbl and MreBH, in cell morphogenesis of Bacillus subtilis. Molecular microbiology. 2009; 73(4):719–731.

Klumpp S, Scott M, Pedersen S, Hwa T. Molecular crowding limits translation and cell growth. Proceedings of the National Academy of Sciences. 2013; 110(42):16754–16759.

Knapp BD, Odermatt R Rojas ER, Cheng W, He X, Huang KC, Chang F. Decoupling of rates of protein synthesis from cell expansion leads to supergrowth. Cell systems. 2019; 9(5):434–445.

Koch AL. The surface stress theory of microbial morphogenesis. Advances in microbial physiology. 1983; 24:301–366.

Konopka MC, Sochacki KA, Bratton BP, Shkel IA, Record MT, Weisshaar JC. Cytoplasmic Protein Mobility in Osmotically Stressed Escherichia coli. Journal of Bacteriology. 2009; 191(1):231–237. https://jb.asm.org/content/191/1/231, doi: 10.1128/JB.00536-08.

Koo BM, Kritikos G, Farelli JD, Todor H, Tong K, Kimsey H, Wapinski I, Galardini M, Cabal A, Peters JM, et al. Construction and analysis of two genome-scale deletion libraries for Bacillus subtilis. Cell systems. 2017; 4(3):291–305.

Marquis RE. Immersion refractometry of isolated bacterial cell walls. Journal of bacteriology. 1973; 116(3):1273–1279.

Matias VR, Beveridge TJ. Cryo-electron microscopy reveals native polymeric cell wall structure in Bacillus subtilis 168 and the existence of a periplasmic space. Molecular microbiology. 2005; 56(1):240–251.

Meeske AJ, Riley EP, Robins WP, Uehara T, Mekalanos JJ, Kahne D, Walker S, Kruse AC, Bernhardt TG, Rudner DZ. SEDS proteins are a widespread family of bacterial cell wall polymerases. Nature. 2016; 537(7622):634–638.

Merchante R, Pooley HM, Karamata D. A periplasm in Bacillus subtilis. Journal of bacteriology. 1995; 177(21):6176–6183.

Mercier R, Kawai Y, Errington J. Excess membrane synthesis drives a primitive mode of cell proliferation. Cell. 2013; 152(5):997–1007.

Miller I, Zsigray R, Landman O. The formation of protoplasts and quasi-spheroplasts in normal and chloramphenicol-pretreated Bacillus subtilis. Microbiology. 1967; 49(3):513–525.

Mindich L. Membrane synthesis in Bacillus subtilis: I. Isolation and properties of strains bearing mutations in glycerol metabolism. Journal of molecular biology. 1970; 49(2):415–432.

Mindich L. Studies on bacterial membrane biogenesis using glycerol auxotrophs. In: Membrane Biogenesis Springer; 1975.p. 429–454.

Misra G, Rojas ER, Gopinathan A, Huang KC. Mechanical consequences of cell-wall turnover in the elongation of a Gram-positive bacterium. Biophysical journal. 2013; 104(11):2342–2352.

Müller A, Wenzel M, Strahl H, Grein F, Saaki TN, Kohl B, Siersma T, Bandow JE, Sahl HG, Schneider T, et al. Daptomycin inhibits cell envelope synthesis by interfering with fluid membrane microdomains. Proceedings of the National Academy of Sciences. 2016; 113(45):E7077–E7086.

Noga MJ, Büke F, van den Broek NJ, Imholz NC, Scherer N, Yang F, Bokinsky G. Posttranslational Control of PlsB Is Sufficient To Coordinate Membrane Synthesis with Growth in Escherichia coli. Mbio. 2020; 11(4).

Oldewurtel ER, Kitahara Y, Cordier B, Özbaykal G, van Teeffelen S. Bacteria control cell volume by coupling cell-surface expansion to dry-mass growth. bioRxiv. 2019; p. 769–786.

Omura S. The antibiotic cerulenin, a novel tool for biochemistry as an inhibitor of fatty acid synthesis. Bacteriological reviews. 1976; 40(3):681.

Patel Y, Zhao H, Helmann JD. A regulatory pathway that selectively up-regulates elongasome function in the absence of class A PBPs. Elife. 2020; 9:e57902.

Paton JC, May BK, Elliott WH. Cerulenin inhibits production of extracellular proteins but not membrane proteins in Bacillus amyloliquefaciens. Microbiology. 1980; 118(1):179–187.

Perlmann GE, Longsworth L. The specific refractive increment of some purified proteins. Journal of the American Chemical Society. 1948; 70(8):2719–2724.

Popham DL, Setlow P. Phenotypes of Bacillus subtilis mutants lacking multiple class A high-molecular-weight penicillin-binding proteins. Journal of Bacteriology. 1996; 178(7):2079–2085.

Pulschen AA, Sastre DE, Machinandiarena F, Crotta Asis A, Albanesi D, de Mendoza D, Gueiros-Filho FJ. The stringent response plays a key role in B acillus subtilis survival of fatty acid starvation. Molecular microbiology. 2017; 103(4):698–712.

Rock CO, Cronan JE. Escherichia coli as a model for the regulation of dissociable (type II) fatty acid biosynthesis. Biochimica et Biophysica Acta (BBA)-Lipids and Lipid Metabolism. 1996; 1302(1):1–16.

Rojas E, Theriot JA, Huang KC. Response of Escherichia coli growth rate to osmotic shock. Proceedings of the National Academy of Sciences. 2014; 111(21):7807–7812.

Rojas ER, Huang KC, Theriot JA. Homeostatic cell growth is accomplished mechanically through membrane tension inhibition of cell-wall synthesis. Cell systems. 2017; 5(6):578–590.

Schindelin J, Arganda-Carreras I, Frise E, Kaynig V, Longair M, Pietzsch T, Preibisch S, Rueden C, Saalfeld S, Schmid B, et al. Fiji: an open-source platform for biological-image analysis. Nature methods. 2012; 9(7):676–682.

Schujman GE, Choi KH, Altabe S, Rock CO, de Mendoza D. Response of Bacillus subtilis to cerulenin and acquisition of resistance. Journal of bacteriology. 2001; 183(10):3032–3040.

Sharpe ME, Hauser PM, Sharpe RG, Errington J. Bacillus subtilis cell cycle as studied by fluorescence microscopy: constancy of cell length at initiation of DNA replication and evidence for active nucleoid partitioning. Journal of bacteriology. 1998; 180(3):547–555.

Sun Y, Garner EC. PrkC modulates MreB fllament density and cellular growth rate by monitoring cell wall precursors. BioRxiv. 2020;.

Theisen A. Refractive increment data-book for polymer and biomolecular scientists. Nottingham University Press; 2000.

Tinevez JY, Perry N, Schindelin J, Hoopes GM, Reynolds GD, Laplantine E, Bednarek SY, Shorte SL, Eliceiri KW. TrackMate: An open and extensible platform for single-particle tracking. Methods. 2017; 115:80–90.

Turner RD, Hurd Ap Cadby A, Hobbs JK Foster SJ. Cell wall elongation mode in Gram-negative bacteria is determined by peptidoglycan architecture. Nature communications. 2013; 4(1):1–8.

Ursell T, Lee TK; Shiomi D, Shi H, Tropini C, Monds RD, Colavin A, Billings G, Bhaya-Grossman I, Broxton M, et al. Rapid, precise quantification of bacterial cellular dimensions across a genomic-scale knockout library. BMC biology. 2017; 15(1):17.

Vadia S, Jessica LT, Lucena R, Yang Z, Kellogg DR, Wang JD, Levin PA. Fatty acid availability sets cell envelope capacity and dictates microbial cell size. Current Biology. 2017; 27(12):1757–1767.

Vazquez A. Optimal cytoplasmatic density and flux balance model under macromolecular crowding effects. Journal of theoretical biology. 2010; 264(2):356–359.

Wang Z, Millet L, Mir M, Ding H, Unarunotai S, Rogers J, Gillette MU, Popescu G. Spatial light interference microscopy (SLIM). Optics express. 2011; 19(2):1016–1026.

Yang D, Männik J, Retterer ST, Männik J. The effects of polydisperse crowders on the compaction of the Escherichia coli nucleoid. Molecular microbiology. 2020; 113(5):1022–1037.

Zhou HX, Rivas G, Minton AP. Macromolecular crowding and confinement: biochemical, biophysical, and potential physiological consequences. Annu Rev Biophys. 2008; 37:375–397.

Zielińska A, Savietto A, de Sousa Borges A, Martinez D, Berbon M, Roelofsen JR, Hartman AM, de Boer R, Van der Klei IJ, Hirsch AK, et al. Flotillin-mediated membrane fluidity controls peptidoglycan synthesis and MreB movement. Elife. 2020; 9:e57179.

Zimmerman SB, Trach SO. Estimation of macromolecule concentrations and excluded volume effects for the cytoplasm of Escherichia coli. Journal of molecular biology. 1991; 222(3):599–620.

